# Inhibiting microglia proliferation after spinal cord injury improves recovery in mice and nonhuman primates

**DOI:** 10.1101/2021.03.06.434049

**Authors:** Gaëtan Poulen, Emilie Aloy, Claire M. Bringuier, Nadine Mestre-Francés, Emaëlle V.F. Artus, Maïda Cardoso, Jean-Christophe Perez, Christophe Goze-Bac, Hassan Boukhaddaoui, Nicolas Lonjon, Yannick N. Gerber, Florence E. Perrin

**Author notes:** Corresponding author: Florence E. Perrin University of Montpellier, MMDN, INSERM Place Eugène Bataillon CC105, 34095 Montpellier Cedex 5 France. These authors contributed equally (Yannick N. Gerber, Florence E. Perrin). The authors have declared that no conflict of interest exists.

## Abstract

No curative treatment is available for any deficits induced by spinal cord injury (SCI). Following injury, microglia undergo highly diverse activation processes, including proliferation, and play a critical role on functional recovery.

In a translational objective, we investigated whether a transient pharmacological reduction of microglia proliferation after injury is beneficial for functional recovery after SCI in mice and nonhuman primates. The colony stimulating factor-1 receptor (CSF1R) regulates proliferation, differentiation, and survival of microglia, we thus used an oral administration of GW2580, a CSF1R inhibitor.

First, transient post-injury GW2580 administration in mice improves motor function recovery, promotes tissues preservation and/or reorganization (identified by coherent anti-stokes Raman scattering microscopy), and modulates glial reactivity.

Second, post-injury GW2580-treatment in nonhuman primates reduces microglia proliferation, improves functional motor function recovery, and promotes tissue protection. Notably, three months after lesion microglia reactivity returned to baseline value.

Finally, to initiate the investigation on molecular mechanisms induced by a transient post-SCI GW2580-treatment, we used microglia-specific transcriptomic analysis in mice. Notably, we detected a downregulation in the expression of inflammatory-associated genes and we identified genes that were up-regulated by SCI and further downregulated by the treatment.

Thus, a transient oral GW2580 treatment post-injury may provide a promising therapeutic strategy for SCI patients and may also be extended to other central nervous system disorders displaying microglia activation.

## Introduction

Traumatic spinal cord injury (SCI) results in 0.6 to 0.9 million annual new cases worldwide [1], induces sensory, motor, and autonomic deficits ranging from minimal dysfunctions to complete tetraplegia. There is no curative treatment available.

Following traumatism, microglia, the resident immunocompetent cells of the central nervous system (CNS) modulate neuroinflammation by releasing both detrimental and beneficial factors to their surrounding cells [for review see [2]]. Microglia response occurs within minutes after SCI and is followed by infiltration of neutrophils and monocyte-derived macrophages from the periphery by 6 hours and 3 days post-lesion, respectively [for review see [3, 4]]. Infiltrating macrophages suppress microglial activation by reducing their expression of inflammatory molecules and ability to phagocytose, consequently preventing chronic microglia-mediated inflammation and blocking these infiltrating macrophages reduces functional recovery after SCI [5]. Notably, microglia exhibit greater SCI-induced proliferation than infiltrating macrophages [6]. Moreover, microglial molecular response after SCI is characterized by an early proliferation followed by a concomitant upregulation of pro- and anti-inflammatory factors [7].

Microglia express the receptor for macrophage colony stimulating factor-1 (CSF1R). CSF1R activation regulates proliferation, differentiation, and survival of myeloid lineage cells. CSF1R inhibition triggers microglial demise *in vitro* and *in vivo* [for review see [8]]. Recent studies have targeted CSF1R to modulate neuroinflammatory response in physiological [9, 10] and multiple neuro-pathological conditions [11–14] [for review see [15, 16]]. PLX5622 is a CSF1R inhibitor that eradicates selectively and almost entirely microglia. Four weeks following traumatic brain injury in mice, a 1-week PLX5622-treatment reduced several histological markers of the injury and was associated with improved motor and cognitive functions recovery [17]. PLX5622 had also been used in mouse models of SCI [18, 19]. Continuous treatment between 3 weeks prior to SCI and 35 days post-lesion worsened motor function, whilst a brief post-injury treatment between 1-6 days transiently deteriorated locomotion up to 21 days after moderate thoracic (T9/T10) contusion [50 kilodynes, kdyn] [18]. PLX5622 treatment for 6 weeks post-SCI, improved cognition and depressive-like behavior recovery but did not improve the overall locomotor activity even if the regularity index and stride length were enhanced at 6 weeks following moderate/severe T10 contusion [60-70kdyn] [19]. PLX3397, another specific CSF1R inhibitor, given to mice between 7 days prior to complete T10 crush injury and 4 weeks after lesion impaired locomotor recovery, disorganized glial scar formation, increased lesion size, and reduced neuronal survival [20].

GW2580, an inhibitor of the tyrosine kinase activity of CSF1R and, to a lesser extent, other related kinases such as FMS tyrosine kinase 3 (FLT3, CD135) and oncogene KIT (c-Kit, CD117), selectively inhibits microglia/monocytes proliferation [21, 22]. GW2580 treatment ameliorates neurological outcomes in animal models of multiple sclerosis [23], prion disease [24, 25], Alzheimer’s disease [26], Parkinson’s disease [27], lupus [28], and amyotrophic lateral sclerosis [29]. Recently, we have shown that continuous GW2580-treatment between 4 weeks prior to a T9 lateral hemisection and up to 6 weeks post-lesion in mice decreases microglia proliferation, reduces gliosis, microcavity formations, and improves fine motor recovery [22]. To enhance translational research, it is critical to examine the effect of GW2580 treatment after SCI not only in mice but also in nonhuman primates. Indeed, pathological responses to SCI vary considerably between rodents and primates due to differences in the neuroanatomical organization of motor and sensory systems, and neurophysiological variations [30, 31].

In the current study, we show in mice that a 1-week GW2580 oral treatment starting immediately after T9 lateral hemisection improves functional recovery, preserves myelin structure, and modifies glial reactivity. We then extend our investigations to *Microcebus Murinus*, a small nonhuman primate and show that orally administered GW2580 over 2 weeks after SCI transiently reduces microglia proliferation and improves motor function recovery. Finally, we used cell-specific transcriptomic analysis in mice to initiate the investigation on molecular mechanisms induced by a transient post-SCI GW2580-treatment. Notably, we identified genes that were up-regulated by SCI and further downregulated by the treatment.

## Results

### Transient post-injury CSF1R blockade in mice enhances motor recovery after SCI

To study the effect of reducing microglia proliferation after SCI, we administered a transient (1 week) oral GW2580 treatment to CX3CR1^+/eGFP^ mice immediately after T9 lateral hemisection. We examined motor recovery over 6 weeks post-injury. GW2580 treatment improved both static and dynamic parameters. The “print position” ipsilateral to the lesion recovered better in the GW2580 group (**Figure 1A**). The “print position” reflects the distance between the front and the hind paws located on the same side in a step cycle and thus measures normality of the hind paw motricity. The “regularity index” measuring inter-paw dynamic coordination also recovered better in the GW2580 group (**Figure 1B**). Indeed, almost full coordination from 3 weeks after SCI was observed in the treated group inversely to the untreated group. The “max intensity” (print maximum intensity) of the hind paw ipsilateral to the lesion (**Figure 1C**) and the “max contact” (max intensity of the paw at max contact) (**Figure 1D**) also recovered better in GW2580-treated mice than in the untreated controls. Improvements appeared at 4 weeks post-lesion. Therefore, a transient 1-week GW2580 treatment immediately after SCI promotes motor function recovery in mice.

**Figure 1:**
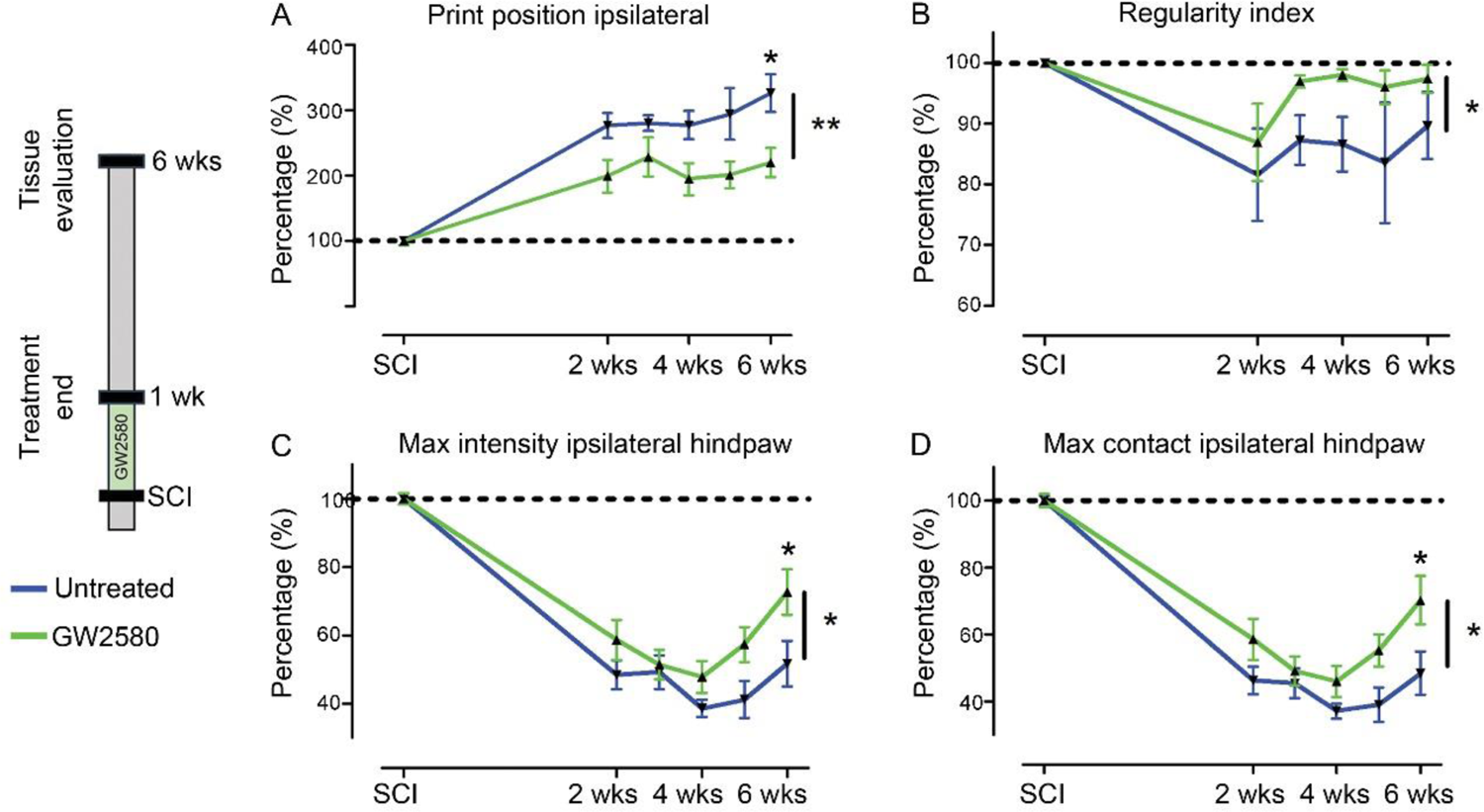
Transient CSF1R blockade after SCI in mice improves motor function recovery. CatWalk^TM^ behavioral analysis (**A-D**). Values were normalized to those obtained by the same animal prior to the lesion (represented as dash lines, 100%). Graphs display in both GW2580-treated and untreated groups the print position of the paw ipsilateral to the lesion (**A**), the regularity index (**B**), the max intensity of the ipsilateral hind paw (**C**), and the maximum intensity at max contact of the ipsilateral hind paw (**D**). In all graphs, results obtained by untreated mice are in blue and GW2580-treated mice in green. Data are mean ± SEM per group. Two-Way ANOVA followed by Bonferroni post-hoc tests, *p < 0.05 and **p < 0.01. Number of mice: n=10 in each group.

### Transient post-injury GW2580 treatment in mice prevents myelin break-down

To investigate whether GW2580 treatment could affect myelin preservation we used coherent anti-stokes Raman scattering (CARS) microscopy (**Figure 2**). Using a scoring method to assess label-free multiphoton myelin morphology [32], we first scored myelin degradation on sagittal spinal cord sections of treated and untreated mice (**Figure 2A-D**) at 6 weeks after SCI. We assessed myelin morphology on 3 locations in close vicinity of the lesion (lesion epicenter and 0.97mm rostral and caudal to the lesion, **Figure 2A**) on the ipsilateral and contralateral sides of the lesion site. Scores ranged from 0 (normal white matter) to 3 (loss of axonal alignment and predominant lipid debris (**Figure 2B**). No significant difference between groups was observed ipsilateral and contralateral to the lesion (**Figure 2C**&**D**). Using axial spinal cord sections (**Figure 2K**&**L**), we then quantified the density of intact myelinated fibers (**Figure 2E-H**, arrows in **E**&**F, I-J** zoom of boxes in **E** and **F**, respectively) at the injury epicenter and more distally from the lesion site (3.15mm rostral and caudal) in both treated and untreated animals at 6 weeks after SCI. Three distinct fields were analyzed in the lateral *funiculi* (**Figure 2K**-**L**) ipsilateral (**Figure 2G**) and contralateral (**Figure 2H**) to the lesion. GW2580 treatment induced a higher intact myelinated fibers density as compared to control ipsilateral to the lesion in the caudal segment (**Figure 2G**) but not contralateral to the lesion side (**Figure 2H**). Altogether, these results demonstrate that a transient GW2580-treatment after SCI in mice decreases myelin break-down following injury.

**Figure 2:**
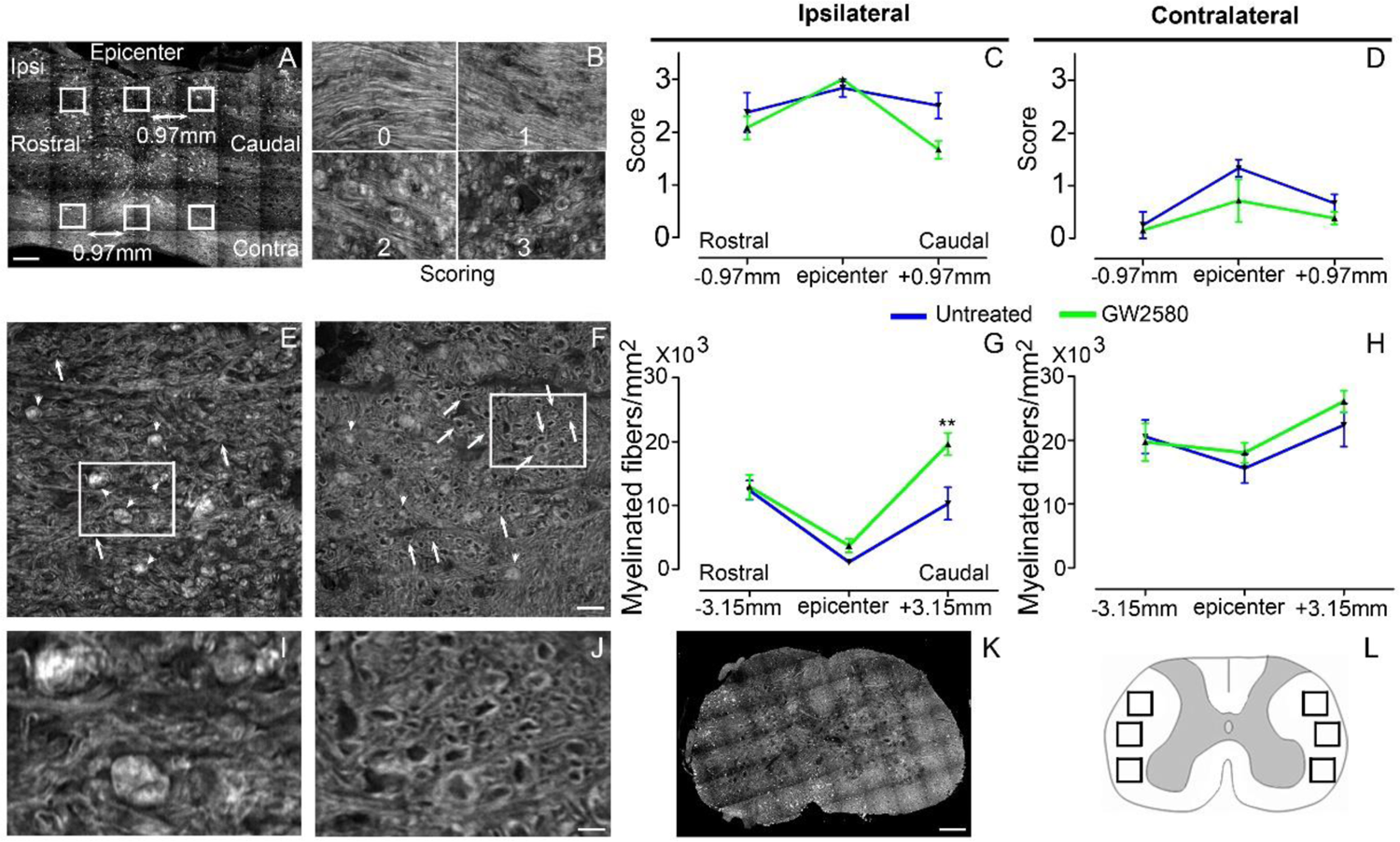
Transient CSF1R blockade after SCI in mice prevents myelin break-down Sagittal CARS low resolution mosaic of a mouse spinal cord to indicate locations of the 6 images acquired per mouse (white boxes) used to score myelin morphology (**A**). Myelin scorings (**B**), normal white matter is associated with the score zero; scores 1 and 2 reflect an increasing occurrence of lipid debris and disorganized axonal arrangement and score 3 represents a complete loss of axonal alignment and major lipid debris. Myelin morphology scores quantified 6 weeks after SCI on sagittal sections of the spinal cord ipsilateral (**C**) and contralateral (**D**) to the lesion site. Representative CARS axial imaging of myelin after SCI in untreated (**E**&**I**) and GW2580-treated (**F**&**J**) mice. Quantification on axial sections of myelinated fibers/mm^2^ ipsilateral (**G**) and contralateral (**H**) to the lesion site 6 weeks after a lateral hemisection of the spinal cord in untreated and treated groups. Axial CARS low resolution mosaic of a mouse spinal cord (**K**). Schematic spinal cord, boxes indicate locations of the 6 images acquired per mouse to quantify myelinated fibers density (**L**). In all graphs, results obtained by untreated mice are in blue and GW2580-treated mice in green. Scale bars: 500µm (**A**&**K**); 20µm (**B**); 20µm (**E-F**), and 5µm (**I**&**J**). Number of mice: n=3 for untreated and treated groups for both axial and sagittal sections. Myelinated fibers: for each animal, 3 images per rostro-caudal location (−3.15mm, epicenter and +3.15mm) were quantified on both the ipsilateral and contralateral sides. Data are mean ± SEM per group. Student’s unpaired t-test, *p < 0.05.

### GW2580 treatment after SCI in mice modulates microglial reactivity

We then quantified microglial reactivity at 2 and 6 weeks after SCI on a 1cm-perilesional segment of the spinal cord of untreated (**Figure 3A**) and GW2580 treated mice (**Figure 3B**); we used IBA1, a specific microglia marker. As we observed a differential microglia activation after SCI within the dorsal *funiculus* as compared to the overall white matter, we analyzed separately the dorsal *funiculus* (**Figure 3C-D**; **G-H** and **K-L**) and the white matter excluding the dorsal *funiculus*, (**Figure 3E-F**&**I-J**). Notably, an analysis of IBA1 expression in the dorsal *funiculus* 6 weeks after SCI along the rostro-caudal axis revealed a higher microglia activation in the rostral segment on both ipsilateral (**Figure 3C**) and contralateral (**Figure 3D**) sides of the spinal cord as well as a constant higher microglia activation in the GW2580 group. Two weeks after lesion, in the white matter of the rostral segment, IBA1 expression was similar between GW2580-treated and untreated groups (**Figure 3I**) conversely to the ipsilateral side of the caudal segment where GW2580 treated mice presented a lower IBA1 expression than the control (**Figure 3J**, p<0.01). In the dorsal *funiculus*, 2 weeks after SCI, IBA1 expression was similar in both segments in the two groups (**Figure 3K-L**). Six weeks after lesion, the white matter of GW2580-treated mice displayed a higher IBA1 expression on the ipsilateral side of the spinal cord in the rostral segment (**Figure 3E-F**&**I**, p<0.01) and on both sides of the spinal cord in the caudal segment (**Figure 3J**, ipsilateral p<0.05, contralateral p<0.01). In the dorsal *funiculus*, IBA1 expression was also higher contralateral to the lesion in the rostral segment (**Figure 3G-H**&**K**), conversely to the caudal segment where microglial reactivity was similar in both groups (**Figure 3L**).

**Figure 3:**
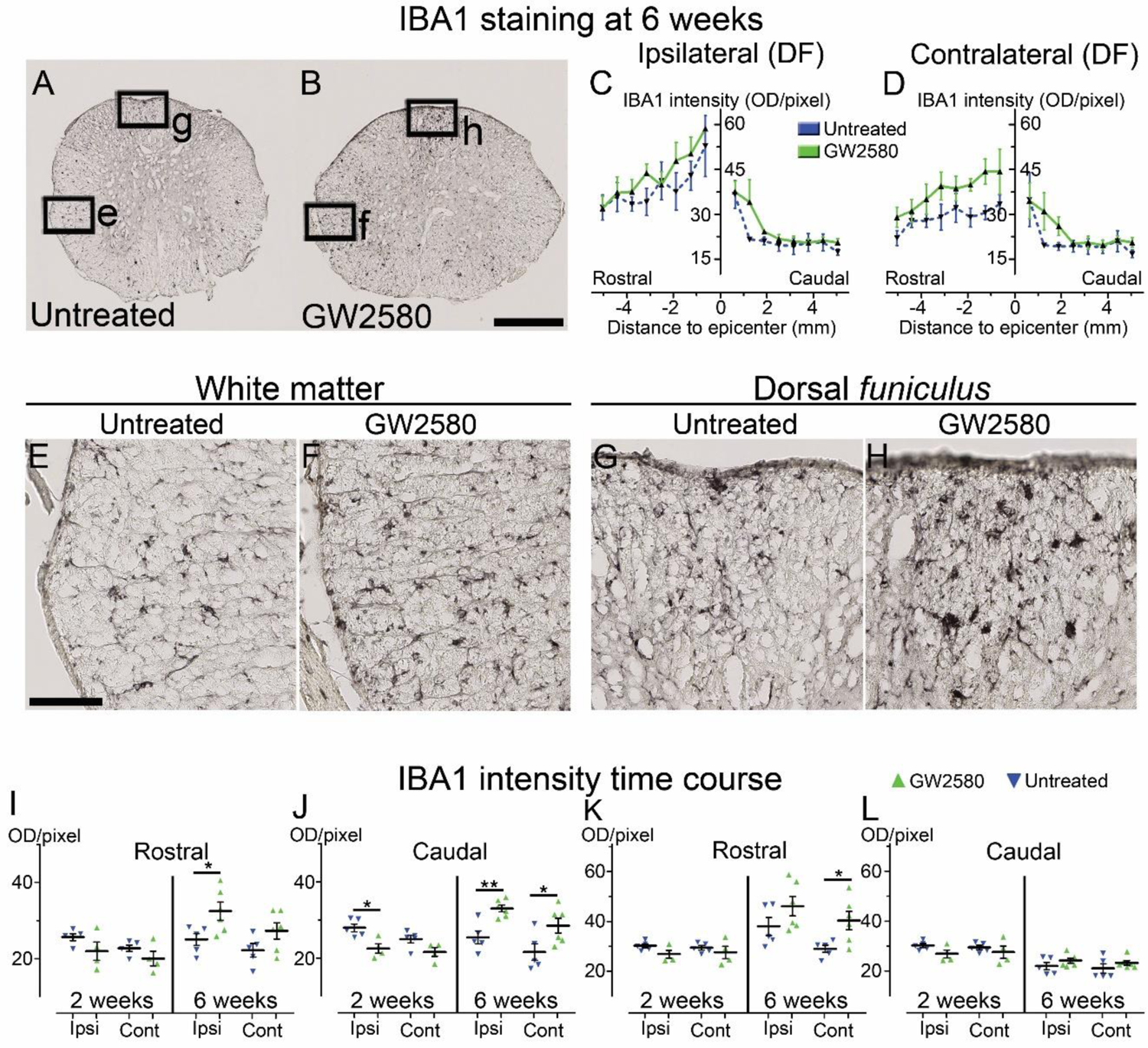
Transient CSF1R blockade after lateral spinal cord hemisection in mice modulates microglial reactivity Bright-field micrographs showing IBA1-positive microglia at 6 weeks after SCI in untreated (**A**, **E**&**G**) and GW2580-treated (**B**, **F**&**H**) mice rostral to the lesion site. Higher magnifications (**E**-**H**) of black insets in **A**&**B**. Line curves display quantification of IBA1-immunoreactivity in the *dorsal funiculus* on the ipsilateral (**C**) and the contralateral (**D**) sides of the injured spinal cord along the rostro-caudal axis. Quantifications of IBA1-immunoreactivity in segments rostral (**I**&**K**) and caudal (**J**&**L**) to the lesion. Quantification in the white matter (excluding the *dorsal funiculus*) (**I**&**J**) and the *dorsal funiculus* (**K**&**L**) at 2 and 6-weeks following SCI. IBA1-immunoreactivity was quantified on ipsilateral and contralateral sides of the spinal cord. Scale bars: 500µm (**A-B**), and 100µm (**E**-**H**). Number of mice: 2 weeks n=5 for untreated and n=4 for treated; 6 weeks n=5 for untreated and n=6 for treated. Data are mean ± SEM per group. Student’s unpaired t-test, *p < 0.05, **p < 0.01.

These data suggest that a transient 1-week inhibition of microglia proliferation in mice using GW2580, slightly reduces IBA1 expression in the white matter 2 weeks after SCI. This is followed by an overall increase in IBA1 expression 6 weeks after lesion.

### Transient GW2580 treatment post-SCI in nonhuman primates reduces microglia proliferation

In order to investigate whether oral GW2580 treatment would also induce beneficial effects on functional recovery after spinal cord injury in nonhuman primates, we first examined whether GW2580 reduces microglia proliferation after lateral hemisection of the spinal cord in *Microcebus murinus* as we previously demonstrated in mice [22]. For *Microcebus murinus*, we chose a dose of 7.2mg/day based on the amount given to mice and a treatment duration of 2 weeks since microglial activation is delayed in nonhuman primates compared to rodents [33]. Two *Microcebus murinus* underwent a lateral hemisection of the spinal cord, one was untreated and the second was treated with 7.2mg/day of GW2580 for 2 weeks. Animals were daily injected with BrdU during treatment and their spinal cords were analyzed. Two weeks following SCI, microglia (IBA1^+^, **Figure 4A**&**E**), proliferating cells (BrdU^+^, **Figure 4B**&**F**) and proliferating microglia (IBA1^+^/BrdU^+^ cells, **Figure 4C**&**G** and **D**-**H**) were stained in untreated (**Figure 4A**-**D**) and GW2580-treated *Microcebus murinus* (**Figure 4E**-**H**). Images were taken contralateral to the lesion site at the epicenter. In a first set of fluorescent images acquisition, we observed an overall decrease in microglia proliferation in the GW2580-treated animal (**Figure 4G**, arrows) as compared to the non-treated one (**Figure 4C**, arrows). We then acquired a second set of images using confocal microscopy to further confirm IBA1^+^/BrdU^+^ colocalization in untreated (**Figure 4D**, arrows) and GW2580-treated animals (**Figure 4H**, arrows). These findings demonstrate that GW2580 treatment in nonhuman primates reduces microglial proliferation after SCI.

**Figure 4:**
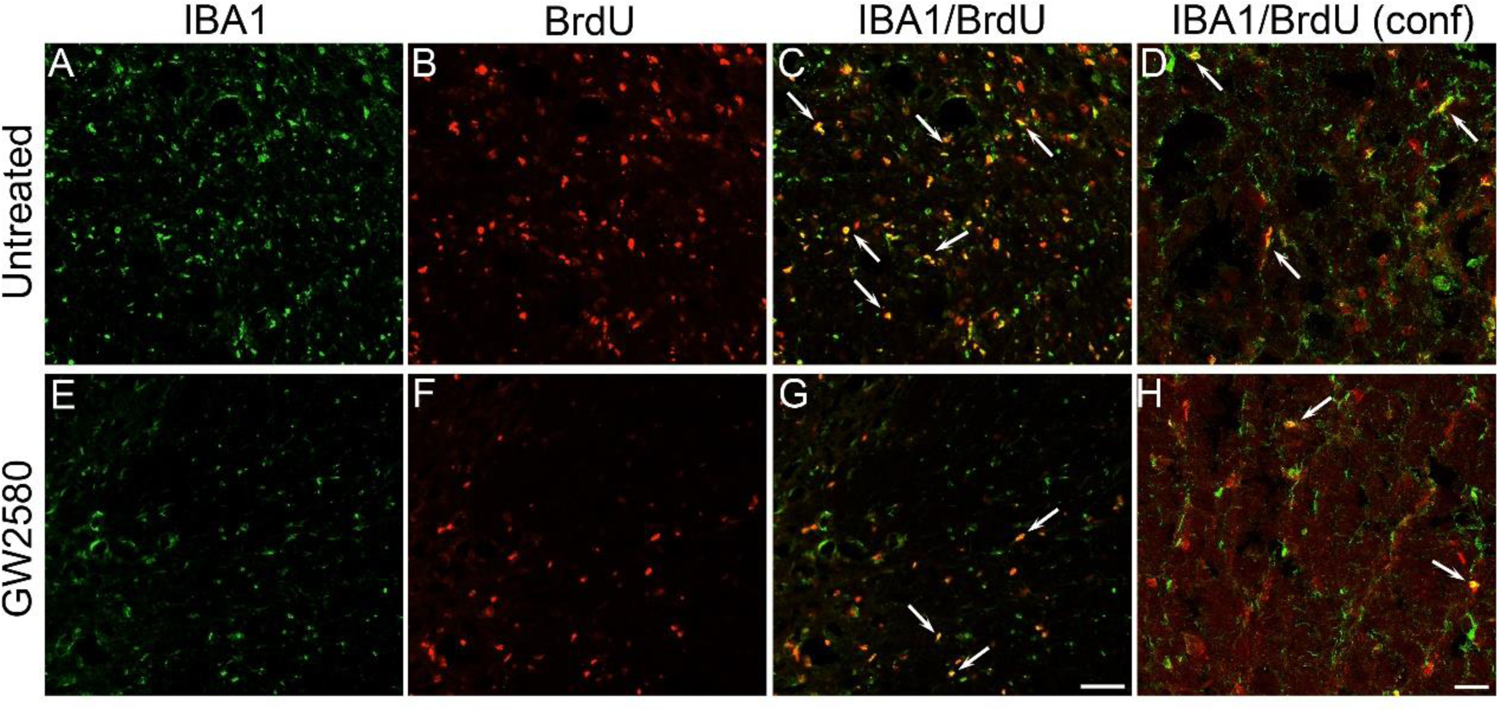
Transient CSF1R blockade after lateral spinal cord hemisection in nonhuman primate decreases microglia proliferation Fluorescent micrographs of axial spinal cord sections from untreated (**A**-**D**) and GW2580-treated (**E**-**H**) *Microcebus murinus* at 2 weeks after SCI. All pictures were taken on the contralateral side at the lesion epicenter. IBA1 staining (**A**&**E**), BrdU staining (**B**&**F**) and merged (**C**&**G**). Confocal microscopy of merged IBA1 and BrdU staining (**D**&**H**). Arrows (**C**&**G** and **D**&**H**) indicate proliferative microglia (BrdU^+^/IBA1^+^). Scale bars: 50µm (**A-C** and **E-G**), and 20µm (**D**&**H**). Number of *Microcebus murinus*: n=1 for untreated and n=1 for treated animals.

### Transient GW2580 treatment post-SCI in nonhuman primates improves motor recovery

We next investigated whether oral GW2580 treatment would also induce beneficial effects on functional recovery after lateral spinal cord hemisection in nonhuman primates. We thus performed a low thoracic (T12-L1) lateral hemisection of the spinal cord in 10 adult males, 5 were treated orally with GW2580, while 5 were untreated. Following SCI, all animals presented a hind limb monoplegia ipsilateral to the spinal cord lesion; only 1 animal developed a transient and incomplete deficit of the contralateral hind limb that spontaneously recovered within 72 hours. All animals survived the surgery, none developed bladder dysfunction, self-biting, cutaneous infection or inflammation and none presented body weight decrease except on days following SCI that is partly due to the 12 hours of fasting prior to anesthesia. Behavioral CatWalk™ assessments were conducted prior to surgery and from 1 day to 3 months after injury (**Figure 5A**). Pre-operative runs were similar in both groups (**Figure 5A**, D0). Immediately after injury, in both groups and for all parameters we observed a severe decrease in performance compared to pre-operative values (100%) (**Figure 5A-E**), as indicated by the absence of the print corresponding to the hind paw located on the ipsilateral side of the lesion (**Figure 5A**, arrows in D0 compared to D1). Motor function displayed a significant improved recovery in the treated group as compared to the untreated, as attested by the re-appearing of a consistent print of the hind paw located on the ipsilateral side of the lesion from 14- and 49-days post-injury in the GW2580-treated and untreated groups, respectively (**Figure 5A**, arrows). We selected 4 accurate criteria to quantify motor function recovery: 2 static parameters i.e. the base of support of the hind paws (distance between hind paws) and the print length of the hind paw ipsilateral to the lesion and 2 dynamic parameters i.e. the regularity index (inter-paw coordination), and the duration of the swing phase (the time without contact of a given paw to the glass plate in a step cycle). The base of support of hind paws returned to pre-operative values (100%) at 14 days post-injury in the treated group, contrariwise to the untreated group that returned to normal value at 80 days after SCI (**Figure 5B**, 2-Way ANOVA **p <0.01). The print length of the ipsilateral hind paw returned to normal from day 10 in GW2580-treated animals contrariwise to the untreated group that did not returned to the pre-operative value though the 3 months post lesion (**Figure 5C**, 2-Way ANOVA **p <0.01). The regularity index displayed an earlier (from 7 days post-injury) and enhanced recovery in the treated group as compared to the untreated group (**Figure 5D**, 2-Way ANOVA *p <0.05). Finally, recovery of the swing of the ipsilateral hind paw also occurred significantly earlier in the GW2580-treated group than in the control group (**Figure 5E**, 2-Way ANOVA *p <0.05). We then developed two additional tests *i.e.* the ladder (**Figure 5F**), and the grip tests (**Figure 5H**) specifically for *Microcebus murinus*. Scoring of the ipsilateral hind paw at the ladder test before injury was 10/10 for all; 24 hours after SCI animals all scored 0/10 (**Figure 5G**). Three months, after SCI treated animals displayed a score of 8/10 while remaining around 6/10 in the untreated group (**Figure 5G**). Similarly, the grip score was also discriminative since treated animals almost returned to a normal pre-operative score (score=4) approximately 35 days after injury conversely to the untreated group that barely returned to normal values (**Figure 5I**). Taken together, these data demonstrate that a transient GW2580-treatment after SCI improves both static and dynamic parameters of motor function recovery in nonhuman primate.

**Figure 5:**
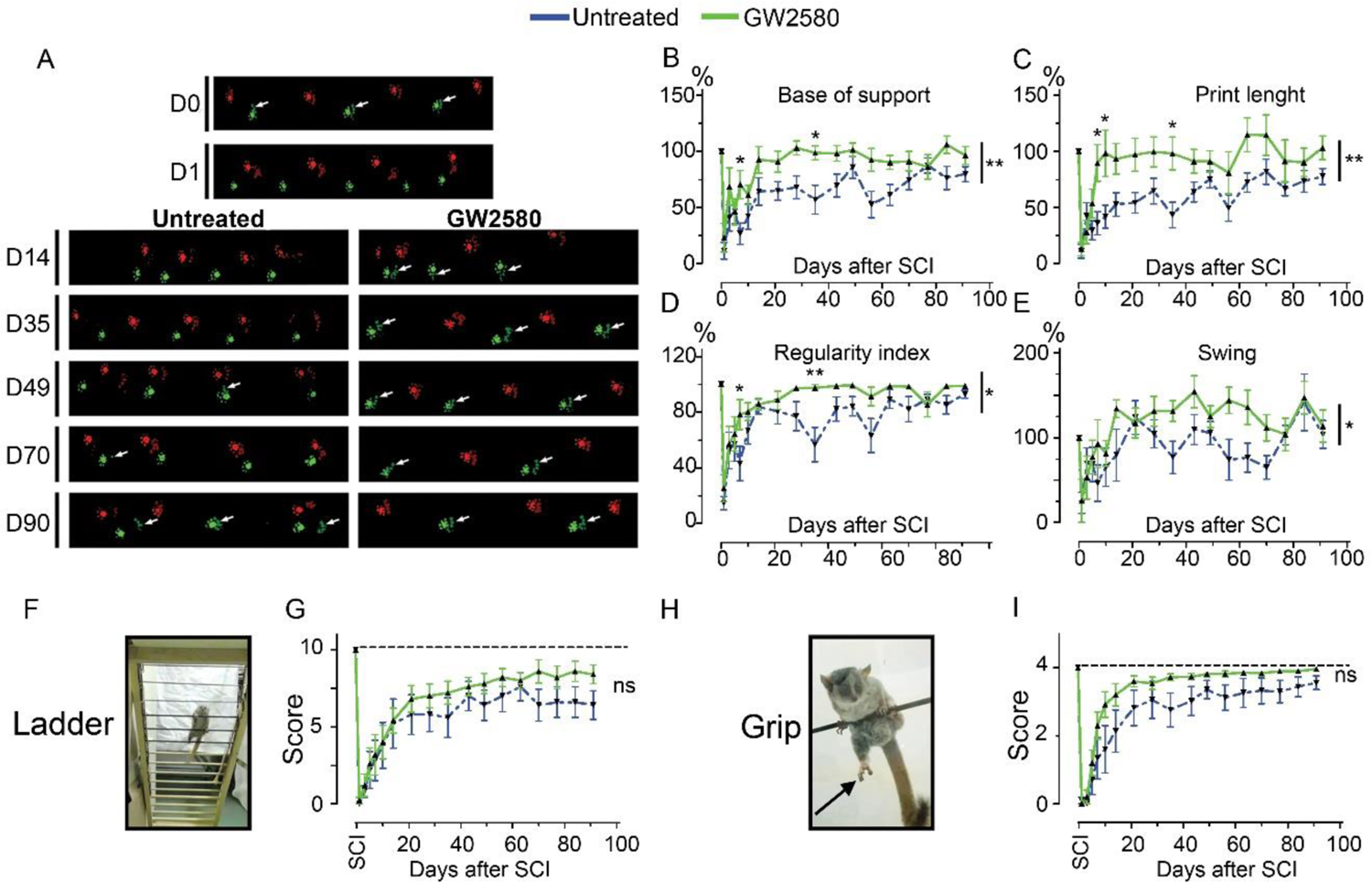
Transient CSF1R blockade after lateral spinal cord hemisection in nonhuman primates improves motor function recovery Representative CatWalk™ runs of *Microcebus murinus* before (D0) and after (D1 to D90) lateral spinal cord hemisection (**A**). Front paws are represented in bright, hind paws in matte, contralateral paws in red and ipsilateral paws in green. White arrows point to the hind limb located on the injured side of the spinal cord. Line graphs displaying the base of support of hind paws (**B**), the print length of the hind paw on the injured side of the spinal cord (**C**), the regularity index (**D**), and swing phase of the hind limb located on the injured side of the spinal cord (**E**). Photographs of the ladder (**F**) and the bar (**H**) tests used to score the grip function of nonhuman primates. Arrow points the hind limb located on the injured side of the spinal cord. Line graphs displaying scores obtained with the ladder (**G**) and grip (**H**) tests. In all graphs, results for untreated nonhuman primates are in blue and GW2580-treated in green. Data are mean ± SEM. Two-Way ANOVA followed by Bonferroni post-hoc tests, *p < 0.05 and **p <0.01. Number of injured *Microcebus murinus*: untreated n=5 and GW2580-treated for 2 weeks n=5.

### Transient GW2580-treatment after SCI in nonhuman primates participates in tissue reorganization and prevents myelin break-down

To study the effect of GW2580 treatment on lesion size and spinal cord microstructure, in particular myelin, we used *ex vivo* diffusion-weighted magnetic resonance imaging (DW-MRI). White and grey matters as well as the lesion site were clearly identifiable (**Figure 6A**-**C**). The lesion epicenter was identified as the section with the highest percentage of damaged tissues (**Figure 6B**, **D**&**E**). Three months after lesion, percentage of damaged tissues at the injury epicenter (50% in both groups), lesion extension (5.6+/-0.5 and 5.4+/-0.4mm in the untreated and GW2580 treated group, respectively) and lesion volume (AUC 168+/-22.5 and 120.3+/-35.6 in the untreated and GW2580 treated group, respectively) were similar in both groups (**Figure 6E**&**F**). Rostral to the lesion, we observed a hypersignal in both sides of the dorsal *funiculus* (DF) in the untreated group and only on the ipsilateral side of the lesion in GW2580-treated animals (**Figure 6G**, red arrows). We thus quantified longitudinal (**Figure 6H-L**) and transverse (**Figure 6M**) diffusivities on a 2 cm-spinal cord segment centered on the lesion site separately in the white matter (WM) excluding the *dorsal funiculus* and the DF. LADC was similar in both the WM and the DF of GW2580-treated as compared to untreated animals rostral and caudal to the lesion (**Figure 6I-J**). However, an analysis of LADC in the WM (without DF, **Figure 6K**) and the DF (**Figure 6L**) along the rostro-caudal axis highlighted that the GW2580-treated LADC curve was always above the non-treated curve. No difference was detected between groups for TADC (data not shown).

**Figure 6:**
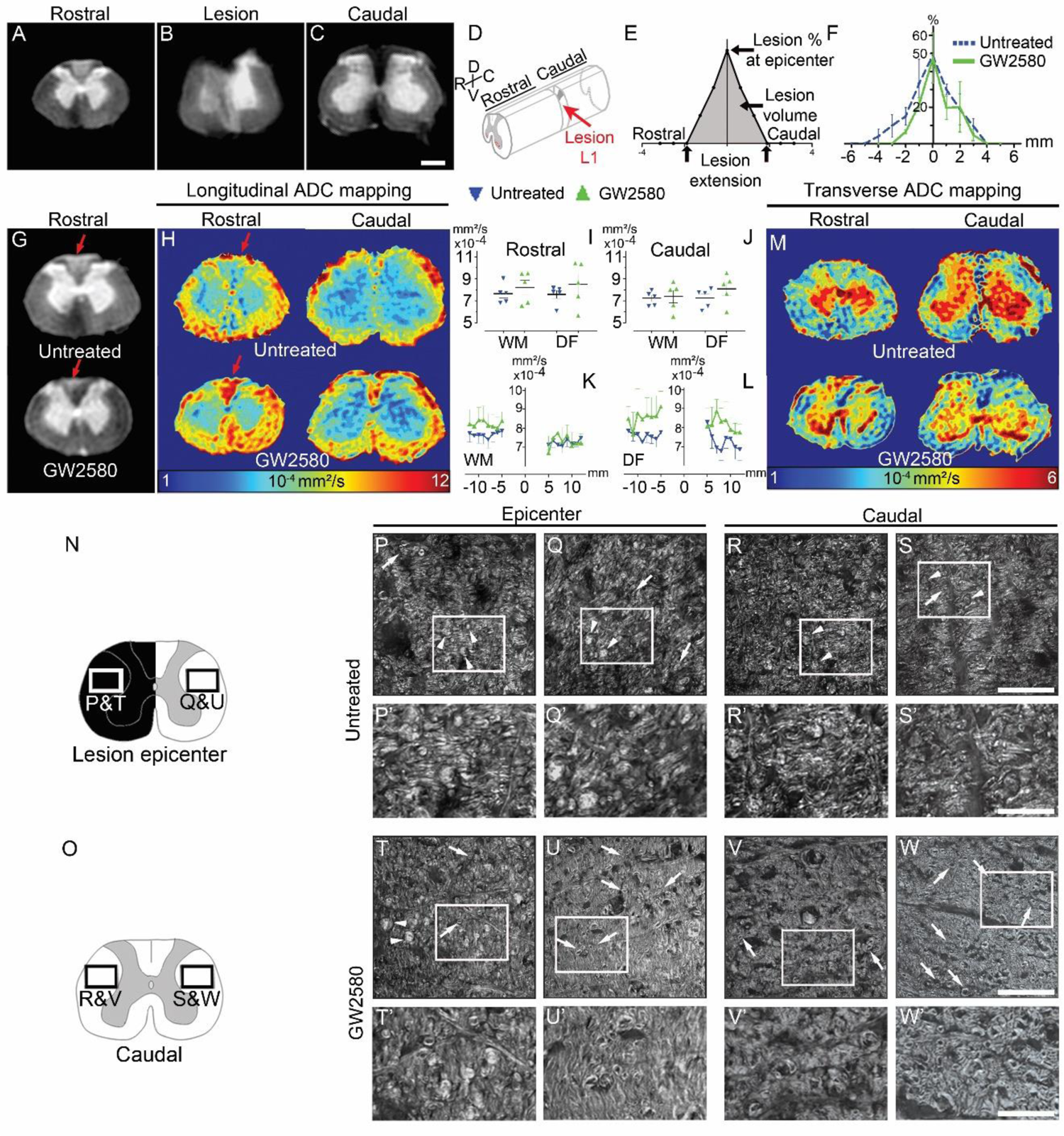
GW2580 treatment after SCI in nonhuman primates preserves white matter ADC and protects myelin *Ex vivo* diffusion-weighted MRI rostral (**A**), within (**B**), and caudal (**C**) to the lesion 3 months after SCI in an untreated lemur. Schematic view of a T12-L1 lateral spinal cord hemisection (**D**). Schematic drawing of quantified parameters (**E**). Quantification 3 months following injury of the lesion percentage at the epicenter, the lesion extension and volume (area under the curve) (**F**). *Ex vivo* DW-MRI (**G**), longitudinal (**H**) and transverse (**M**) ADC mapping in treated and untreated animals. Red arrows in (**G&H**) indicate hyper-intense signal on both sides of the dorsal *funiculus* (DF) (untreated) and only on the hemisected side (GW2580). Longitudinal (**I**-**J**) diffusivities in the white matter and the DF. Quantifications were done rostral (**I**) and caudal (**J**) to the lesion epicenter. Quantification of LADC in the white matter (without DF) (**K**) and the DF (**L**) along the rostro-caudal axis. Schemes of the spinal cord at the lesion epicenter (**N**) and 2.1mm caudal to the lesion (**O**). CARS imaging (**P**-**W’**) taken in insets presented in **N**&**O**. Myelin organization after SCI in untreated (**P**-**S’**) and GW2580-treated (**T**-**W’**) primates at the lesion epicenter (**P**&**P’**; **Q**&**Q’**; **T**&**T’** and **U-U’**) and caudal (**R**&**R’**; **S**&**S’**; **V**&**V’** and **W**&**W’**) to the lesion. Images ipsilateral (**P**&**P’**, **T**&**T’**, **R**&**R’**; and **V**&**V’**) and contralateral (**Q**&**Q’**; **U**&**U’**, **S**&**S’**; and **W**&**W’**) to the lesion. Insets in **P**-**S** and **T**-**W** correspond to higher magnifications in **P’**-**S’** and **T’**-**W’** respectively. Results for untreated nonhuman primates are in blue and GW2580-treated in green. Data are mean ± SEM per group. Student’s unpaired t-test, *p < 0.05. Scale bars (**A-C&G**): 600µm; (**P**-**S** and **T**-**W**): 50µm and (**P’**-**S’** and **T’**-**W’**): 20µm. Number of animals for MRI experiments: 5 untreated and 5 GW2580-treated and 1 animal in each group for CARS experiments.

We further assessed the effect of GW2580-treatment on SCI-induced demyelination using Luxol fast blue and neutral red staining on the same spinal cord 1-cm segment (**Supplementary Figure 1**). Quantification of myelin damage in the spinal cord using Luxol highlighted a reduction, that however did not reach statistical significance, in the lesion percentage at the injury epicenter in the GW2580-treated group (44.45±8.7 and 29.15±4.8% in untreated and treated groups, respectively, **Supplementary Figure 1C-D**). No difference between groups in the spared white matter rostral (**Supplementary Figure 1A-B**) and caudal (**Supplementary Figure 1E-F**) to the lesion epicenter was seen on both ipsilateral and contralateral sides. To further investigate if GW2580 treatment could act on the overall protection of myelin break-down we used CARS on axial sections of the spinal cord of untreated (**Figure 6P-S’**) and GW2580-treated animals (**Figure 6T-W’**) taken at the lesion epicenter (**Figure 6N**, **P-Q**; **P’-Q’**, **T-U** and **T’-U’**) and 2.1mm caudal to the lesion (**Figure 6O**, **R-S**; **R’-S’**, **V-W** and **V’-W’**) on both ipsilateral (**Figure 6P**&**P’**and **T**&**T’**; **R**&**R’** and **V**&**V’**) and contralateral sides (**Figure 6Q**&**Q’**and **U**&**U’**; **S**&**S’**and **W**&**W’**) of the lesion. Preserved myelin sheaths were almost absent in all sections of untreated animals (**Figure 6P-S**, arrows and **P’-S’**). Conversely, well distinguishable intact myelin sheaths were seen in the contralateral side of the spinal cord of treated animals (**Figure 6U**&**U’**, and **W**&**W**’, arrows) and to a lesser extent on the ipsilateral side of the lesion (**Figure 6T**&**T’** and **V**&**V**’, arrows). Lipid accumulation, most likely formed by myelin debris, were also regularly observed in the spinal cord of untreated *Microcebus murinus* (**Figure 6P**&**Q** and **P’**&**Q’**; **R**&**S** and **R’**&**S’** arrowheads) contrariwise to GW2580-treated animals (**Figure 6T**&**U** and **T’**&**U’**; **V**&**W** and **V’**&**W’** arrowheads). Thus, a transient GW2580 treatment post-SCI in nonhuman primates evokes an overall protection of white matter degradation (*ex vivo* DW-MRI) and an overall protection of myelin breakdown (CARS).

### Microglia reactivity returns to baseline 3 months after GW2580 treatment in nonhuman primates

Similarly, to mice we quantified microglial reactivity at 3 months after SCI on a 1cm-perilesional segment of the spinal cord using IBA1 (**Figure 7**). We did not observe a differential activation of microglial cells in between untreated (**Figure 7A, E**&**G**) and GW2580-treated nonhuman primates (**Figure 7B, F**&**H**), neither within the white matter (excluding the dorsal *funiculus*) (**Figure 7I**&**J**) nor in the dorsal *funiculus* (**Figure 7K**&**L**). However, in the dorsal *funiculus* rostral to the lesion, microglial reactivity was overall limited to the ipsilateral side of the lesion in GW2580-treated nonhuman primates (**Figure 7B**, **C**&**D**, **H**) while it usually spreads in the entire *funiculus* in the untreated group (**Figure 7A**, **C**&**D, G**). These data suggest that a transient inhibition of microglia proliferation using GW2580, does not affects overall IBA1 expression in the long term after SCI in nonhuman primates.

**Figure 7:**
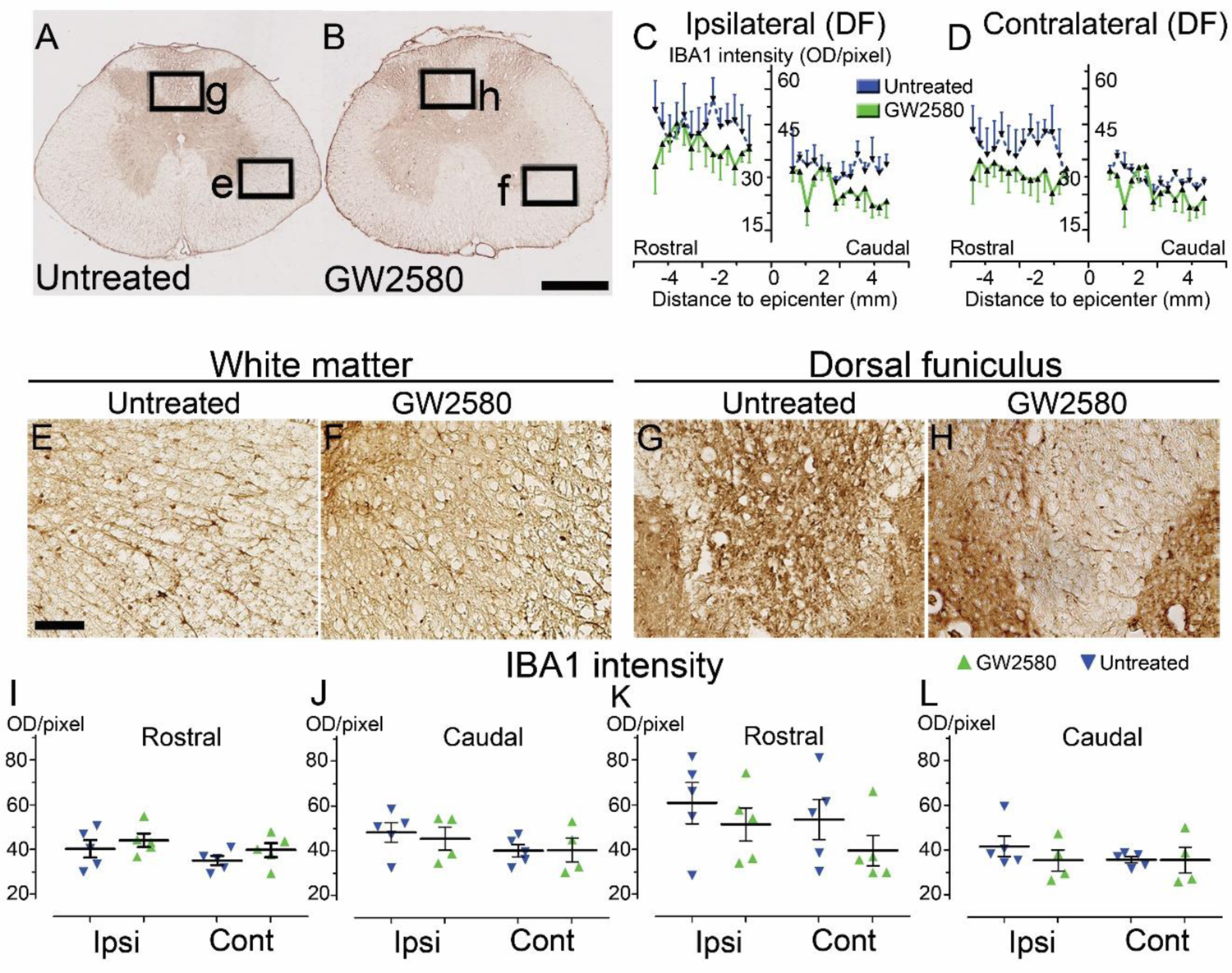
Transient CSF1R blockade after lateral spinal cord hemisection in nonhuman primates does not affect microglial reactivity in the long term Bright-field micrographs showing IBA1-positive microglia after SCI in untreated (**A**, **E**&**G**) and GW2580-treated (**B**, **F**&**H**) nonhuman primates rostral to the lesion site 3 months after SCI. Higher magnifications (**E**-**H**) of black insets in **A**&**B**. Line curves display quantification of IBA1-immunoreactivity in the dorsal *funiculus* on the ipsilateral (**C**) and the contralateral (**D**) sides of the injured spinal cord along the rostro-caudal axis. Quantifications of IBA1-immunoreactivity in segments rostral (**I**&**K**) and caudal (**J**&**L**) to the lesion. Quantification in the white matter (excluding the dorsal *funiculus*) (**I**&**J**) and the dorsal *funiculus* (**K**&**L**) at 3-months following SCI. IBA1-immunoreactivity was quantified on ipsilateral and contralateral sides of the spinal cord (**I**-**L**). Scale bars (**A**&**B**): 500µm; (**E**&**H**): 100µm. At least 40 sections (centered on the lesion site) per animal at 210µm intervals were analyzed. Number of *Microcebus murinus*: injured & untreated n=5, injured & GW2580-treated for 2 weeks n=5.

### GW2580 treatment after SCI does not modify muscle fibers surface and neuromuscular junction density in nonhuman primates

Improvement in motor function recovery following SCI induced by GW2580 treatment suggests a better preservation of the skeletal hind limb muscles. We thus analyzed in both groups muscle fiber surface and the neuromuscular junctions (NMJ) density in the *gastrocnemius* of both hind limbs (**Figure 8A-D**). Overall, muscle fiber surface (**Figure 8E**) and NMJ density (**Figure 8F**) were similar in *gastrocnemius* located on the ipsilateral and the contralateral sides of the spinal cord lesion in both untreated and treated groups. Comparison between groups did not identify significant difference for both parameters. Therefore, GW2580 does not induce quantifiable difference between the ipsilateral and the contralateral muscle fiber surface and NMJ density of *gastrocnemius* three months after SCI in nonhuman primates.

**Figure 8:**
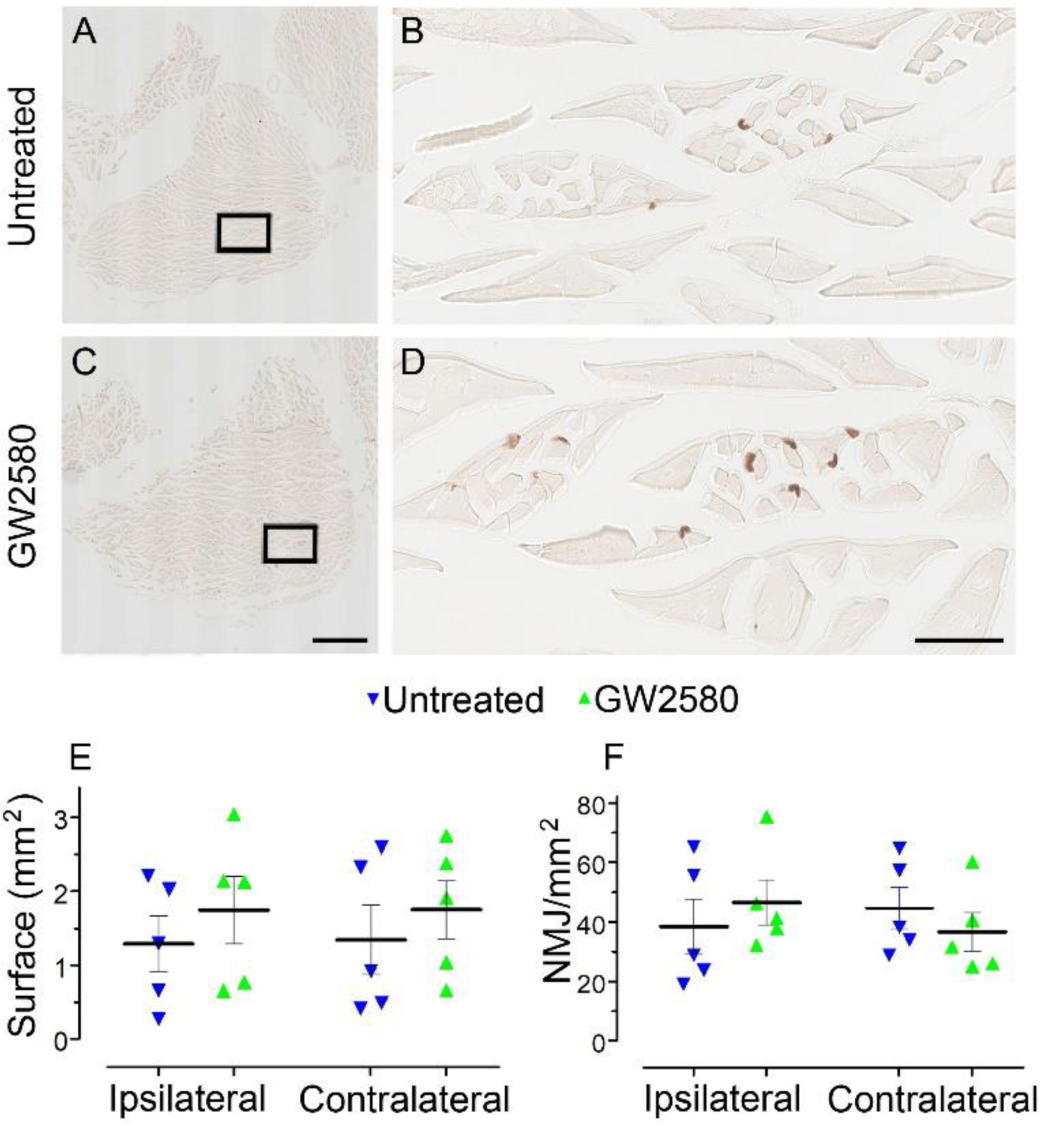
Transient CSF1R blockade after lateral spinal cord hemisection in nonhuman primates does not affect muscle surface and neuromuscular junction density Bright-field micrographs showing *gastrocnemius–soleus–plantaris* muscle complex of the hind limb located on the ipsilateral side of the spinal cord lesion in untreated (**A**) and GW2580-treated (**C**) *Microcebus murinus*. Black boxes in **A**&**C** correspond to higher magnification taken within the *gastrocnemius* muscle and presented in **B**&**D**, respectively. Graphs displaying quantitative assessments of the *gastrocnemius* muscle fiber surface area (**E**) and the density of neuromuscular junctions (**F**). In all graphs, results for untreated nonhuman primates are in blue and GW2580-treated are in green. Data are mean ± SEM per group. Student’s unpaired t-test was used. Scale bars (**A**&**C**): 1mm, (**B**&**D**): 100µm. At least 20 sections per animal throughout the *gastrocnemius* muscle at 16 µm intervals were analyzed. Number of *Microcebus murinus*: injured & untreated n=5, injured & GW2580-treated for 2 weeks n=5.

### Identification by cell-specific transcriptomics of genes up-regulated by SCI and further downregulated by GW2580-treatment

Finally, to initiate investigations on molecular mechanisms induced by a transient post-SCI GW2580-treatment we took advantage of the CX3CR1^+/eGFP^ transgenic mice that express eGFP in microglia. We combined fluorescence-activated cell sorting and RNA-Seq analysis. We isolated only eGFP^high^ positive cells from treated and untreated CX3CR1^+/eGFP^ spinal cord injured mice. Indeed, we have previously shown in this injury model that 1-week after lesion eGFP^high^ expressing cells located in a 1 cm segment centered on the injury site are CD11^+^/LY6C^neg/low^/GFP^high^ and thus mostly correspond to microglia without contamination of infiltrating monocytes [7]. Transcriptomic analyses were done at the end of the treatment (1 week after injury) on microglia from pooled 1cm-spinal cord segments centered on the lesion site (at least 2 animals and 16.000 cells per replicate), as previously described [7]. We selected the same stringent cutoff as in our previous study [Fold Change (FC)≥2 and p-value with false discovery rate (FDR) ≤0.05] [7]. We found 19 differentially expressed (DE) genes; 16 and 3 genes were down and up-regulated, respectively in the GW2580-treated groups as compared to the untreated control (**Figure 9A**-**B** and **Supplementary Table 1**). Thus, microglia from depleted animals clearly displayed a higher number of down than upregulated genes. Unbiased hierarchical clustering revealed a strong reproducibility in DE genes among independent biological replicates (**Figure 9B**). Notably, down regulated genes belong to processes such as regulation of cell proliferation and cell migration [*chondroitin sulfate proteoglycan 4* (*cspg4*), *macrophage scavenger receptor 1* (*msr1*), *glycoprotein transmembrane* (*Gpnmb*), *adrenomedullin* (*adm*) and *fibronectin 1* (*Fn1*)], inflammatory response [*platelet factor 4* (*pf4*), *cytochrome b-245* (*Cybb*)], immune response [*lysozyme* (*Lyz1 and 2*), *cd40* (*tnfrsf5*), *chemokine (C-X-C motif) ligand 13* (*cxcl13*)] (**Supplementary Table 1**). Remarkably, Gene Ontology (GO) enrichment analysis of downregulated genes ranked as top molecular function “CXCR3 chemokine receptor binding (2 DE genes; p-value with FDR: 7.314E-04) and ranked as top processes the inflammatory response (8 DE genes; p-value with FDR 5.588 10E-05, **Supplementary table 2**).

**Figure 9:**
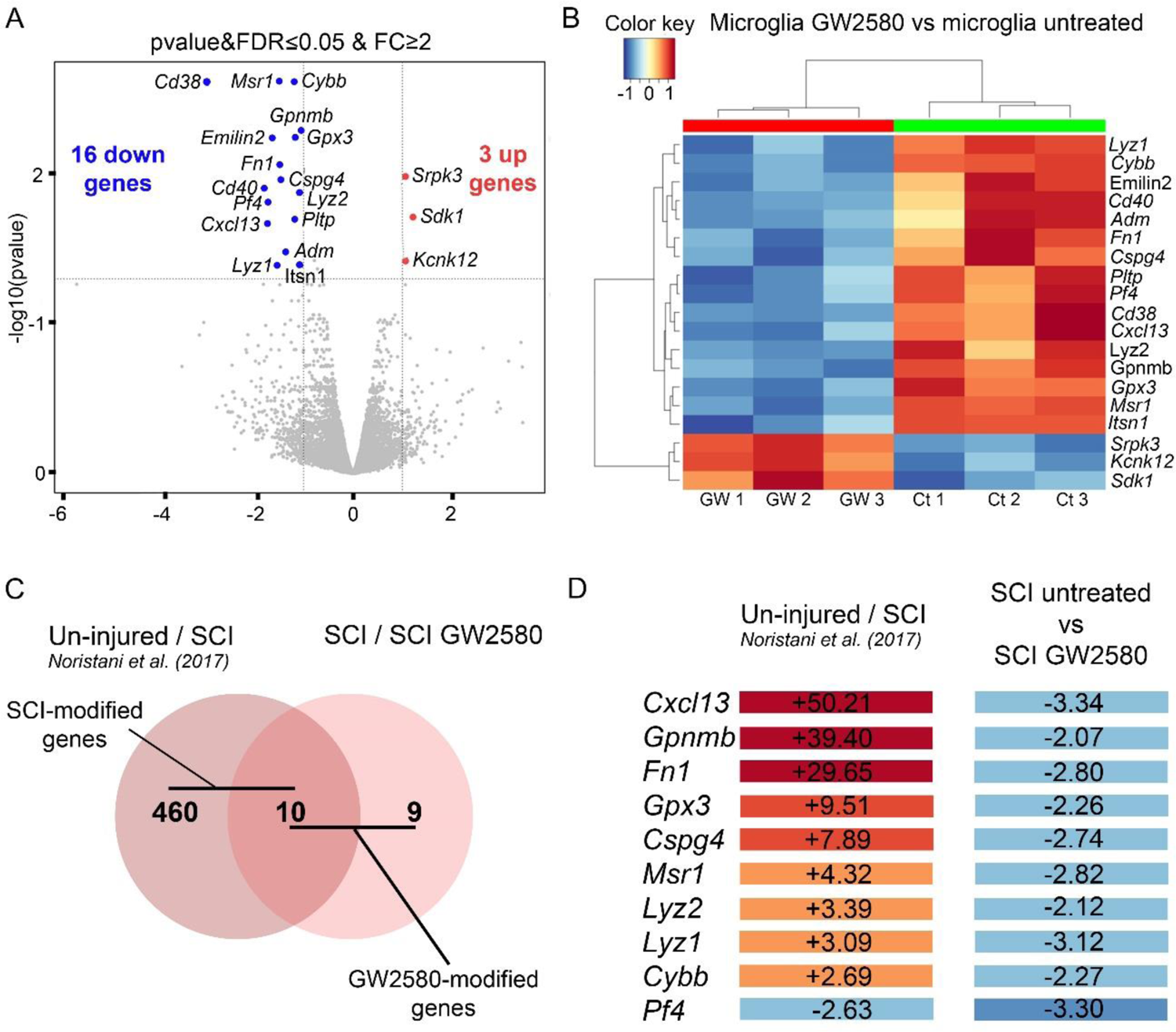
Transient 1-week CSF1R blockade after SCI in mice induces transcriptional modification in microglia RNA-seq analysis of FACS-isolated microglia from pooled (at least 2 animals and 16.000 cells) 1cm-spinal cord segments centered on the lesion site of injured untreated and treated mice at 1 week after injury (end of the treatment) (**A**-**B**). We selected the same stringent cutoff as in our previous study [Fold Change (FC)≥2 and p-value with false discovery rate (FDR) ≤0.05] [7]. Volcano plot (**A**). Heat map (**B**). *In silico* differential expression analysis (**C**-**D**). Comparison of the list of genes deregulated by the injury [identified in our previous study using the same parameters: male CX3CR1^+/eGFP^ mice aged of 3 months, hemisection of the spinal cord at T9 level, analysis of DE genes in microglia in a 1-cm segment centered on the lesion 1 week after injury (uninjured/SCI) [7]] with the list of genes deregulated by GW2580 treatment in SC-injured mice (SCI/untreated *vs* SCI/GW2580). Venn diagram (**C**). Fold changes of the 10 genes commonly deregulated in the comparison between (un-injured/SCI) and SCI-untreated/ SCI GW2580) (**D**).

In a previous study we have identified DE genes in microglia in a 1-cm segment surrounding a T9 spinal cord lateral hemisection in CX3CR1^+/eGFP^ mice 1 week after injury [7]. To determine whether DE SCI-induced genes were modified by GW2580 treatment, we compared the list of genes deregulated by the injury (uninjured *vs* SCI) [7] with the list of genes deregulated by GW2580 treatment in SC-injured mice (SCI/untreated *vs* SCI/GW2580) (**Figure 9C-D**). Out of the 470 DE genes by SCI in microglia, 10 were changed by GW2580. Ninety percent (9 out of 10) of these genes were down-regulated by GW2580 treatment. Strikingly, *cxcl1*3 the most up-regulated gene by SCI (+50.21) is involved in immune response process and 4 out of these 9 genes were involved in cell proliferation and cell migration (*cspg4*, *Gpnmb, Msr1* and *Fn1*). Taken together, these data demonstrate that a transient GW2580 treatment reverse some transcriptional changes in microglial gene expression.

## Discussion

Microglia and macrophages are recognized as pivotal players in SCI pathophysiology [4, 18, 34–36]. They play dual beneficial and detrimental roles after SCI that most likely depend on the kinetics of their responses, including proliferation [3, 4, 18]. Our findings suggest that a transient depletion of microglia proliferation immediately after SCI improves functional recovery in mice and nonhuman primates.

Following SCI, unspecific modulation of anti- or pro-inflammatory responses were often ineffective, if not detrimental. Modulating microglia should thus be done with caution to limit additional risks including induction of systemic diseases already triggered by the SCI-induced disruption of the neural–immune communication [for review see [34]]. We have therefore chosen to transiently inhibit microglial proliferation.

Firstly, we highlighted that a transient GW2580 treatment initiated immediately after SCI enhanced motor recovery in mice and nonhuman primates. Interestingly, improved motor function recovery seems greater in nonhuman primates than in rodents. Enhanced recovery in GW2580-treated lemur started as early as 10-15 days after injury and several parameters eventually returned to pre-injury values. This is consistent with a previous study that also reported better functional recovery in primates than rodents after a lateralized SCI [31]. Besides, we observed in the white matter a trend of higher DW-MRI diffusivities along the rostro-caudal axis in treated, as compared to untreated, nonhuman primates that may attest of tissue reorganization and could reflect the better myelin preservation in treated *Microcebus murinus* 3 months after injury [for review see [37, 38]]. Even if yet not frequently used in SCI [39, 40], Raman spectroscopy can accurately detect SCI-associated microglial inflammation and myelin degradation [32, 41]. Indeed, combination of endogenous two-photon fluorescence and CARS imaging highlighted the presence of lipid debris in SCI-induced inflammatory region thus providing an indicator of activated microglia/macrophages following engulfment of myelin debris [42]. In agreement, the transitory inhibition of microglia proliferation also led to a decreased presence of lipid debris in mice and nonhuman primate and a higher density of myelinated axons in mice 6 weeks after injury.

Our results support recent findings showing that haploinsufficiency of sorting nexin 27 (Snx27) in mice, an endosome-associated cargo adaptor, that amongst other functions suppress microglia/macrophages proliferation, improved motor recovery following SCI [43]. Snx27 deficiency additionally reduces apoptotic neuronal death. Our findings also complement previous work that targeted microglia/monocytes in the context of SCI. First, a selective depletion of a subset of infiltrating Ly6C^+^(Gr1^+^)CCR2^+^ monocytes deteriorated motor recovery following SCI in mice [44]. Second, using CCR2 null mice, it had been shown that stopping the crosstalk between resident microglia and monocyte derived macrophages participates in long-term microglial activation, greater myelin loss that eventually worsen motor function after SCI [5]. Third, complete microglia depletion altered glial scar formation, decreased immune infiltration, neuronal and oligodendrocytes survival associated with impaired motor recovery following SCI in mice [18, 20]. This difference as compared to our findings may result from the specific subpopulation of microglia (*i.e.* only proliferative microglia) that is inhibited by GW2580 [21, 22]. CSF1R is principally expressed by microglia in the intact CNS [45–47]. Therefore, we cannot exclude that GW2580 treatment may likewise have off-target effects and could affect neurons that also express CSF1R [48] or infiltrating macrophages [for review see [15]]. However, SCI induces a sevenfold greater microglia proliferation as compared to infiltrating macrophages [5]

*In vivo,* evidence of a neuroprotective role of proliferating microglia that may serve as an endogenous pool of neurotrophic molecules such as IGF-1 had been reported in cerebral ischemia [49]. The difference may result from the heterogeneity and region-specific differences in microglial response [50]. Greenhalgh et al. highlighted an important phagocytic role of microglia in their early response following SCI associated with a lower susceptibility to cell death associated with a higher microglia proliferation [6] that is consistent with an early upregulation of genes involved in proliferation that peak at 3 days following SCI [7].

In mice, transient inhibition of microglia proliferation induced a reduction of IBA1 expression 2 weeks after SCI followed by an overall increase in its expression 6 weeks after lesion that may correspond to a transient microglial repopulation. Indeed, in *Microcebus murinus*, microglia reactivity returned to baseline 3 months after lesion. Renewal of microglia is observed in physiological condition in mice [51–53] and humans [54]. In pathological conditions, a clonal microglial expansion is reported in mice [52, 53] and in *Macaca fuscata* [55]. Understanding the exact kinetic and extend of microglia proliferation following SCI would raise the possibility of modulating their proliferation in more chronic phases.

Finally, our transcriptomic analyses of microglia highlighted that post-injury GW2580-treatment down-regulated the expression of genes involved in cell proliferation, cell migration and inflammatory response which is consistent with an inhibition of microglia proliferation/activation. We also emphasized that GW2580-treatment reverse the up-regulation of SCI-induced 9 genes and further decreased the expression of 1 SCI-down regulated gene. Interestingly, *cxcl1*3 that is involved in immune response process was strongly up-regulated by SCI (+50.21) and decreased by GW2580 (−3.34), suggesting that GW2580 may inhibit neuroinflammation through CXCL13-mediated signal pathway in SCI. This is consistent with recently reporting in rat that increased miR-186-5p expression following spinal cord ischemia/reperfusion improved neurological outcomes through inhibition of inflammation via several pathways including CXCL13 [56].

Additionally, 4 out of these 9 genes whose expression was reverse by GW2580 are involved in cell proliferation and cell migration that is strongly suggesting inhibition of SCI-induced microglia proliferation/activation. Further investigations of the role of these genes in microglia proliferation will help to better understand molecular mechanisms induced by GW2580-treatment.

*In conclusion,* we show that a transient post-injury oral administration of GW2580, inhibiting microglia proliferation, promotes motor functional recovery and modulates tissue structure following SCI in rodents and nonhuman primates. Beneficial effects of GW2580-treatment on motor recovery seem greater in *Microcebus murinus* than in mice, pointing to the key role of nonhuman primates as critical SCI models to further promote translational research.

## Methods

### Animals

#### Mice

CX3CR1^+/eGFP^ transgenic mice express enhanced green fluorescent protein (eGFP) downstream of the *Cx3cr1* promoter. CX3CR1 is expressed in resident CNS microglia and circulating peripheral monocytes. Mice were maintained on a C57BL/6 background (The Jackson Laboratory, Bar Harbor, ME, USA) and housed in controlled environment (hygrometry, temperature and 12 hours light/dark cycle). Three months old heterozygote males (transcriptomics) and females CX3CR1^+/GFP^ were used.

Number of mice that underwent behavioral tests for 6 weeks: 10 controls and 10 GW2580-treated; 6 mice of each groups also underwent histological analysis. Out of these 6 mice per group, 3 in each group were also analyzed by CARS (myelin). 6 additional mice (3 in each group, 6 weeks post-injury) were dedicated to longitudinal CARS acquisition.

Five controls and 4 GW2580-treated mice were sacrificed at 2 weeks and were investigated by histology. Finally, 16 additional mice (8 controls and 8 GW2580-treated) were devoted to transcriptomic analysis.

#### Nonhuman primates

ten adult males *Microcebus murinus* (2 years old) underwent the complete study (SCI and follow-up over 3 months post-lesion) and 2 injured animals (females, 2 years old) were dedicated to assess GW2580-treatment effects on microglia proliferation. They were all born and bred in the animal facility (CECEMA, University of Montpellier, France) and housed separately in cages (60cm x 60cm x 50cm, equipped with wooden nests and enriched environment) during the entire experiment. Temperature of the animal facility was constantly kept between 24-26°C with 55% humidity. All *Microcebus murinus* were fed 3 times a week with fresh fruits and a mixture of cereal, milk, and eggs. Water was given *ad libitum*. One day prior and after general anesthesia, animals were given mealworms to increase their protein intake.

### Spinal cord injury and post-operative cares

#### Mice

anesthesia was induced with 3-4% isoflurane (Vetflurane®, Virbac, France) and then maintained with a mixture of 1-2 % isoflurane and 1 L/min oxygen flow rate throughout the surgery. Eye gel was applied to the cornea during the surgery. A vertebral laminectomy at thoracic 9 level (T9) followed by a lateral spinal cord hemisection (HS) was done using a micro knife [10315-12, Fine Science Tools (FST)], as described previously [57]. Animals were monitored over 1-hour following the surgery before returning to their cages. Bladders were emptied manually twice daily until recovery of full sphincter control. Bodyweights were monitored prior to surgery and then daily throughout the study. Animals were kept for 2- or 6-weeks post-lesion.

#### Nonhuman primates

Food and water were withdrawn 12 hours prior to surgery. Lateral HS of the spinal cord at low thoracic level (T12 - L1) was done as previously described [58]. Anesthesia was induced with 3-5% isoflurane, eye gel was applied, and anesthesia was maintained with a mixture of 1-2.5 % isoflurane and 1-3 L/min O2 during the surgery. The skin was shaved and cleaned (Vetadine®, Bayer, Australia), cutaneous incision started rostrally at the costovertebral joint of floating ribs and faced 2 vertebral segments. Overlying muscles were disinserted from the midline, a laminectomy followed by a lateral spinal cord HS were done under a microscope (micro knife 10315-12, FST). Lesion quality was assessed by 2 operators and surgical area was profusely cleaned with physiological serum. Muscles and skin were sutured (vicryl, 3/0 and Filapeau 4/0, Braun, Germany, respectively). *Microcebus murinus* were placed on a temperature-controlled pad and monitored over 2 hours before returning to their cages. Animals were observed twice daily to identify potential signs of pain or distress (refusal to eat or drink, absence or decrease in grooming activity, bent posture, self-mutilation). Bladder function was controlled daily. Animals received buprenorphine (0.01mg/kg/day) and amoxicillin (10 mg/kg/jour) for 48 hours after SCI. Animals were kept for 2 weeks (GW2580 effects assessments) or 3 months post-lesion, bodyweights were monitored daily for 2 weeks and then weekly throughout the study.

### Treatment

#### Mice

GW2580 treatment started immediately after the injury and ended 1-week post-lesion. Mice were fed either with a standard rodent chow (A04, maintenance diet, SAFE diet, AUJY, France) or with the same diet containing 0.1% GW2580 (corresponding to 150mg/kg per day per animal, LC Laboratories, Woburn, USA), as previously described [22].

#### Nonhuman primates

we adjusted GW2580 dose based on body surface area (BSA) of animals [59]. BSA of mice and *Microcebus murinus* are approximately 0.007 and 0.016m^2^, respectively. Thus 150mg/kg/day (450mg/m^2^/day) for mice corresponds to approximately 7.2mg/day for *Microcebus murinus*. Six animals received 7.2mg/day of GW2580 and 6 were not treated, it includes 2 animals for GW2580 effects assessments on microglia proliferation. GW2580 treatment started immediately after SCI and ended 2 weeks post-lesion. The treatment was administered *per os*, mixed in a small quantity of applesauce; animals were monitored to ensure that they had taken the treatment.

#### Bromodeoxyuridine (BrdU) experiment

To evaluate GW2580 acute effect on microglia/monocytes proliferation in injured *Microcebus*, we used 2 females (2 years old), both were injured and received a daily injection of BrdU for 2 weeks starting immediately after SCI (s.c., 100mg/kg, Sigma Aldrich, Gilligham, UK) in sterile saline [60], 1 animal was treated for 2 weeks after SCI with GW2580. Animals were sacrificed 24 hours after the last BrdU injection and immunohistochemical analysis for BrdU and IBA1 were done at the lesion site.

### Behavioral assessments

#### Mice

assessments were done at 2 weeks, 1 week and 1 day prior to injury followed by 3 days, 5 days and then once a week up to 6 weeks after lesion (n=10 for untreated and GW2580 groups). Dynamic walking pattern (CatWalk™, XT Noldus, Wageningen, The Netherlands) analysis assessments were done as earlier reported [22, 57, 61]. Six CatWalk™ runs per animal were analyzed per time point. CatWalk™ data analyses were done using CatMerge (Innovationet, Tiranges, France).

#### Nonhuman primates

1-month prior to surgery, habituation to manipulation by the operators and behavioral tests were done 3 times per week to acquire accurate pre-operative values. After injury, tests were done at 1, 3, 5, 7, 10- and 14-days post-surgery and then once a week until 3 months after lesion. Three tests to evaluate the gait and motor activity were done. CatWalk™ Noldus, Wageningen, The Netherlands): 6 runs per animal with at least 3 un-interrupted step cycles were acquired, as previously described [58]. For each animal, values obtained following lesion were normalized to those obtained prior to surgery (results are expressed as percentage of the median pre-operative value). To better assess fine motor recovery, we designed a ladder made of wooden bars with 4 different diameters (15, 10, 5 and 3mm, bottom to top, **Figure 6F**). Animals were placed on the bottom of the ladder and video recorded while climbing toward their nest located at the top of the ladder. Two parameters were quantified for each paw to grade movements (overall movement of a given hind limb and capability to grip) on a scale of 5 and then summed to obtain a value reaching a maximum of 10 in case of normal movements (**Table 1**). To better assess balance and grip recovery, we used a metal bar (diameter: 3mm) located at 20 cm from the ground within an empty Plexiglas test arena (**Figure 5H**). Primates were placed on the bar, and experimenters gently rotated the bar to evaluate the capacity of the animal to use its hind paw and to grip the bar. Animals were video recorded during the whole procedure. Each animal was scored through a motor grading of each hind paw. Final score resulted from the sum of the scores of the overall capacity for the hind paw ipsilateral to the lesion to move toward the bar (0=no movement, 1=incomplete movement, 2=complete movement) and to grip the bar (0=no grip, 1= incomplete grip, 2= complete grip). Video recordings were acquired with a high-resolution camera (HD 1080P, Logitech, Newark, CA, USA) and blindly analyzed by two independent experimenters.

**Table 1.**
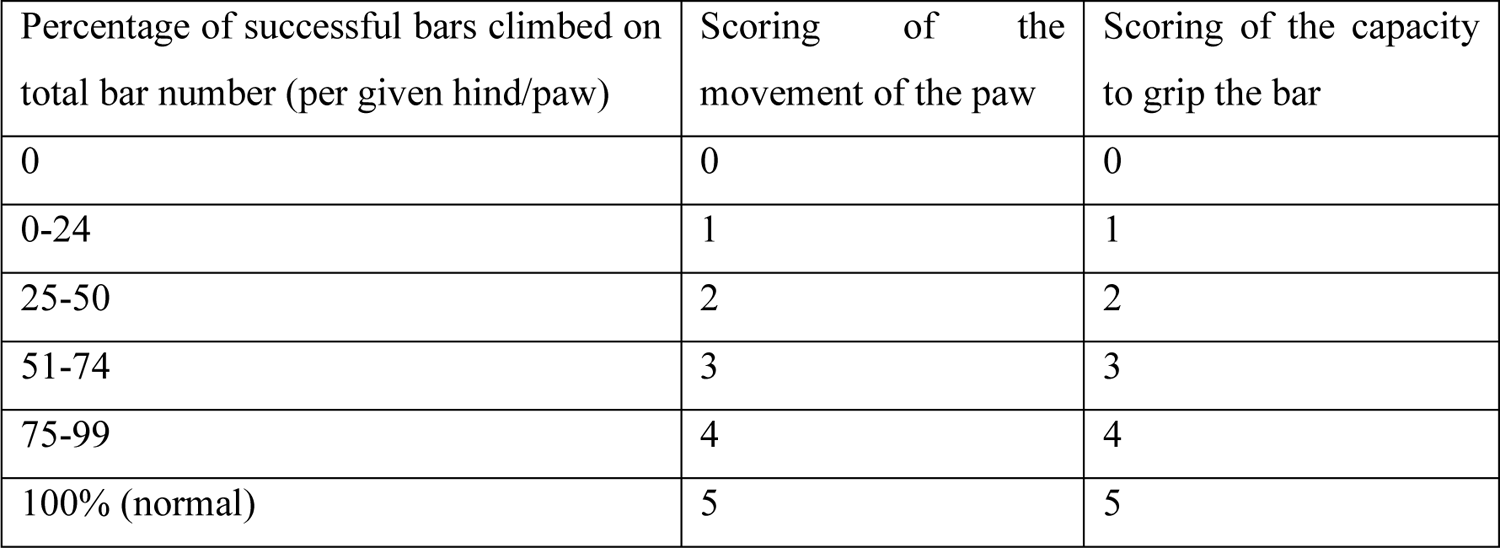
Scoring of the ladder test for *Microcebus murinus*

### *Ex vivo* diffusion-weighted magnetic resonance imaging (DW-MRI)

At 3 months post-injury for nonhuman primates, animals were injected with a lethal dose of ketamine (150 mg/kg, Merial, Lyon, France). Animals were then perfused intracardially with cold phosphate saline buffer (PBS, 0.1M, pH 7.2) followed by cold 4% paraformaldehyde (PFA, pH7.2, Sigma Aldrich, Darmstadt, Germany) in 0.1M PBS. Spinal cords were then dissected and further post-fixed in the same fixative for 2 additional hours and then stored in 1% PFA until *ex vivo* MRI acquisition.

For *ex vivo* acquisition, spinal cords were placed in Fluorinert FC-770 liquid (3M™ Electronic Liquids, Saint Paul, USA) in a 4-mm-diameter glass tube surrounded by a custom-made solenoid coil dedicated to SCI investigations [61, 62]. The coil was placed in the 9.4 Tesla apparatus (Agilent Varian 9.4/160/ASR, Santa Clara, California, USA) associated with a VnmrJ Imaging acquisition system (Agilent, USA). Axial *ex vivo* MRI scans were done using Single Echo Multi Slices (SEMS) sequence (TR=1580ms; TE=30.55ms; AVG=30; FOV=10mm*10mm; 36 slices; thickness=1mm; gap=0mm; acquisition matrix (NREAD*NPHASE)=128*128). Diffusion gradients were applied in 3 directions including the rostro-caudal axis and 2 directions perpendicular to the spinal cord (Gs = 10G/cm; delta = 6.844ms; separation = 15.05ms; b-value= 499.21s/mm²). The same images were also acquired without applying diffusion gradient (Gs=0G/m^-1^). MRI visualizations and segmentations were done manually using Myrian Software (Intrasense, Montpellier, France), as described previously [58]. Longitudinal (LADC) and transversal apparent diffusion coefficients (TADC) were measured on a 2 cm segment centered on the lesion site. Immediately after MRI acquisitions, spinal cords were rinsed in 0.1M PBS, cryoprotected in 30% sucrose, embedded in Tissue Tek (Sakura, Alphen aan den Rijn, The Netherlands), frozen and kept at −20°C.

### Histology

Mice were injected with a lethal dose of tribromoethanol (500 mg/kg, Sigma-Aldrich Darmstadt, Germany) and perfused intracardially and their spinal cord processed and frozen as described above for primates. Serial 14-µm-thick axial spinal cord cryosections (Microm HM550, Thermofisher Scientific, Waltham, USA) were collected on Superfrost Plus© slides.

#### Luxol fast blue and neutral red staining in nonhuman primates

Luxol fast blue staining was done as previously described [63]. Briefly, sections were placed 5 min in 95% ethanol and then incubated in 0.1% Luxol fast blue under mild shaking [12 hours, room temperature (RT)]. Slides were rinsed in milli-Q water (1min), then placed in lithium carbonate (1 min, 0.05%) and finally washed in tap water (1 min). Subsequently, slides were incubated for 10 min in 0.5% neutral red solution, 5 min in 100% ethanol and washed twice for 10 min in xylene. All slides were coverslipped using Eukitt (Sigma Aldrich, Darmstadt, Germany). Quantifications of lesion extension and volume were done on a 1-cm segment centered on the lesion site; sections were analyzed at 210µm intervals in nonhuman primates, respectively. The lesion area was expressed as a percentage of the total surface area; spared white and grey matters were measured (mm^2^).

#### Immunohistochemistry in mice and nonhuman primates

Transversal 14-µm-thick axial spinal cord sections were washed twice in 0.1M PBS, treated for 10 min in 0.1M PBS containing 20mM lysine (pH 7.2) and 15 min in hydrogen peroxide (1% in 0.1M PBS, Sigma Aldrich, Gilligham, UK). Sections were washed twice in 0.1M PBS and blocked for 2 hours with 0.1M PBS containing 1% bovine serum albumin and Triton X-100 (0.1%) (both from Sigma Aldrich, Gilligham, UK) and then incubated for 48 hours at 4°C with the primary antibody, excepted for negative controls. Sections were rinsed with 0.1M PBS (30 min) and incubated in 1:500 dilution of the corresponding biotinylated secondary antibody (2 hours, RT). Sections were rinsed with 0.1M PBS (30 min). For amplification, Avidin Biotin Complex solution (Vector Laboratories Ltd. Peterborugh, UK) diluted at 1:100 in 0.1M PBS was added on slides and incubated (1 hour, RT). Then, sections were rinsed in 0.1M Tris (pH 7.6, RT). Protein expression was visualized using DAB peroxidase substrate kit (Vector Labs, Burlingame, USA). The reaction was stopped by rinsing the sections in 0.1M Tris (3×10 min). Slides were dehydrated in increasing concentrations of ethanol and then xylene. Coverslips were applied using Eukitt (Sigma Aldrich, Darmstadt, Germany). For BrdU detection, sections were first pre-incubated in 2N HCl (hydrogen chloride, 30 min) for DNA (deoxyribonucleic acid) denaturation followed by 0.1M pH 8.5 sodium borate buffer washes (Sigma Aldrich, Gilligham, UK) (3×10 min). Sections were then incubated with a combination of rabbit anti-IBA1and rat anti-BrdU antibodies and then rinsed with PBS (3×10 min). Sections were then incubated in a solution containing the corresponding anti-rat fluorescent and anti-rabbit biotinylated secondary antibodies. Second, a streptavidin fluorescent conjugated antibody was used to amplify IBA1 immunodetection. Sections were cover slipped using fluorescence mounting medium (Dako, Glostrup, Denmark).

#### Antibodies

Primary antibodies: anti-ionized calcium-binding adapter molecule 1 (IBA1, peroxidase staining, 1:1000 for mice, 1:200 for *Microcebus*, Wako Pure Chemical Industries, Japan) and anti-BrdU antibody (1:500; Abcam, Cambridge, UK).

Secondary antibodies: Biotinylated anti-rabbit (1:500, Invitrogen, Carlsbad, USA). Fluorescent anti-rat (Alexa 594) and biotinylated anti-rabbit coupled with streptavidin fluorescent conjugated antibody (Alexa 488) (1:1000, both Life Technologies, Carlsbad, USA).

### Microscopy and quantifications

#### Brightfield microscopy

We used NanoZoomer RS slide scanner with constant light intensity and exposure time and NanoZoomer Digital Pathology System view software (Hamamatsu, Hamamatsu, Japan). To quantify SCI and GW2580 treatment-induced changes in IBA1 expression, the mean optical density (OD) was measured along the spinal cord, as previously described (ImageJ, National Institutes of Health, USA) [22, 57, 58, 61]. To minimize bias in staining intensity, all immunostainings for a given antibody and a given time-point were done in parallel. For all antibodies used, expression levels were analyzed in at least 40 and 16 axial sections throughout the lesion segment of the spinal cord at 210µm and 630µm intervals for *Microcebus Murinus* and mice respectively. OD quantifications included grey and white matters (excluding the dorsal *funiculus*) and dorsal *funiculus*. Background OD was subtracted from OD values of each section. All quantifications were done blindly.

#### Fluorescent microscopies

images were obtained using the Axio Imager 1 microscope (Zeiss, Oberkochen, Germany) and confocal images were obtained with a laser scanning confocal microscope (Leica SPE, Mannheim, Germany) associated with a Leica LAS AF interface. Settings were kept constant for all acquisitions.

#### Coherent anti-stokes Raman scattering (CARS)

We used LSM 7 MP optical parametric oscillator (OPO) multiphoton microscope (Zeiss, Oberkochen, Germany) with an upright Axio Examiner Z.1 optical microscope associated with a femtosecond Ti: sapphire laser (680–1080 nm,80 MHz, 140 fs, Chameleon Ultra II, Coherent, France) pumping a tunable OPOs (1000–1500 nm, 80 MHz, 200 fs, Chameleon Compact OPO, Coherent, France) to acquire CARS images. We imaged axial spinal cord sections (14µm) and longitudinal sections (22µm) at 6 weeks after injury in mice. Axial sections (14µm thickness) were taken at 3 months after injury in primates. A x20 water immersion lens (W Plan Apochromat DIC VIS-IR) with the following characteristics was used: 1024×1024 pixels frame size, scan speed of 6 (zoom x1.2) and 8 (mosaic, zoom x3, PixelDwell 3.15 and 1.27 μs/scan, respectively) and either a zoom x 1.2 or x 3. CARS excites the CH2 vibrational mode at 2845cm^-1^ and CH2 bonds are found in lipids [64]. Excitation wavelengths are 836 and 1097nm (synchronized Ti-saphire and OPO, respectively) and the signal is detected at 675nm (filter from 660-685nm). Pictures are a stack of 3µm (3 slices). Myelin degradation and myelin density were scored and manually quantified, respectively.

#### Neuromuscular junction labelling

Following animal’s perfusion, *gastrocnemius-soleus-plantaris* muscular complex were collected for neuromuscular junction labelling using the method described by Karnovsky and Roots [65]. Axial sections (16µm) of the entire muscle complex were analyzed. Every 15 sections in the *gastrocnemius*, muscle fibers were manually segmented by a blinded experimenter, their surface were quantified and NMJs number was counted.

#### Statistics

Statistical tests were done using GraphPad Prism (GraphPad software 5.03, USA). Significance was accepted at p ≤ 0.05. Results are expressed as mean ± standard error of the mean (SEM). For behavioral analysis, 2-Way ANOVA with Bonferroni post-hoc tests were used. For all other analysis student’s unpaired t-test was used.

#### Transcriptomic analyses

We used Fluorescence Activated Cell Sorting (FACS) to isolate eGFP+ microglia from a 1cm-spinal cord segment centered on the lesion site, as previously described (Noristani, Gerber et al. 2017). Briefly, treated and untreated spinal cord injured CX3CR1^+/GFP^ mice were anesthetized (tribromoethanol, 500mg/kg) and intracardially perfused with 0.1M RNAse-free phosphate base saline (PBS, Invitrogen, Carlsbad, USA). Spinal cords were dissociated in an enzymatic cocktail [750µl PBS, 100µl of 13mg/ml trypsin, 100µl of 7mg/ml hyaluronidase, 50µl of 4mg/ml kinurenic acid (all from Sigma Aldrich, Saint Louis, USA) and 20µl of 10mg/ml DNAse I (Roche, Rotkreuz, Switzerland)] for 30 min at 37°C. Cell suspension was separated (40µm sieve, BD Biosciences, Franklin Lakes, USA), re-suspended in 0.9M sucrose and centrifuged (20min, 750g). Pellet was re-suspended in 500µl of 7AAD 1µl/ml (Sigma Aldrich, Saint Louis, USA) and eGFPhigh expressing cells, that we have previously shown to correspond to microglia (Noristani, Gerber et al. 2017), were sorted using FACS ARIA (BD Biosciences, Franklin Lakes, USA), equipped with a 488nm Laser Sapphire 488–20. Size threshold was used to eliminate cellular debris. Sorted microglia were centrifuged (5min, 700g) and re-suspended in 250µl of RLT lysis buffer (Qiagen, Maryland, USA) and 1% beta-mercaptoethanol. RNA was isolated using the RNeasy Mini Kit, (Qiagen, Maryland, USA, with DNAse) and its quality assessed (Agilent 2100 bioanalyzer, RNA 6000 Pico LabChip, Palo Alto, USA). Only RNA with a RNA integrity number (RIN)>7 were further processed. RNA-Seq was performed on 3 biological replicates per condition (each replicate for both untreated and treated conditions consisted on pooled 1-cm spinal cord segments from at least 2 animals representing a minimum of 16.000 microglia). For reverse transcription and cDNA amplification, we used the SMART-Seq v4 kit (Takara Bio USA, Mountain View, CA, USA) according to manufacturer’s specifications, starting with 2ng of total RNA as input. 200pg of cDNA were used for library preparation using the Nextera XT kit (Illumina, San Diego, CA, USA). Library molarity and quality was assessed with the Qubit and Tapestation using a DNA High sensitivity chip (Agilent Technologies, Santa Clara, CA, USA). Libraries were pooled and loaded for clustering on 1 lane of a Single-read Illumina Flow cell (for an average of 50 million of reads per library) (Illumina, San Diego, CA, USA). Reads of 100 bases were generated using the TruSeq SBS chemistry on an Illumina HiSeq 4000 sequencer (Illumina, San Diego, CA, USA). FastQC was used to assess sequencing quality control, the reads were mapped with the STAR aligner and biological quality control and summarization were done with PicardTools. The counts were produced from aligned reads by the Python software htseq-count with the reference .gtf file. https://htseq.readthedocs.io/en/release_0.11.1/count.html. Normalization and differential expression analysis were performed with the R package edgeR, for the genes annotated in the reference genome http://www.ncbi.nlm.nih.gov/pmc/articles/PMC2796818/. Very lowly expressed genes were filter out to keep only genes that were sufficiently expressed (above 10 in the 3 biological replicates); filtered data were normalized according to the RNA composition and library size. Differentially expressed genes were estimated using a GLM approach (General Linear Model), negative binomial distribution and a quasi-likelihood F-test. Differentially expressed transcripts were defined with a criterion of a 2-fold and greater difference plus a significant p-value ≤ 0.05 with false discovery rate (FDR). Pathway analysis was done using MetaCore, Clarivate Analytic, Philadelphia, USA.

#### Study approval

Experiments were approved by the Veterinary Services Department of Hérault, the regional ethic committee n°36 for animal experimentation, and the French Ministry of National Education, Higher Education and Research (authorizations; mice: n°34118 and nonhuman primates n° APAFIS#16177-2018071810113615v3). Experimental procedures followed the European legislative, administrative and statutory measures for animal experimentation (EU/Directive/2010/63 of the European Parliament and Council) and the ARRIVE guidelines.

## Author contribution

GP performed experiments in nonhuman primates, analyzed the data and contributed to the writing of the manuscript; EA participated in experiments in nonhuman primates and analyzed the data; CMB participated in acquisition and analysis of confocal and MRI experiments; NMF provided lemurs and her expertise in *Microcebus murinus* behavior and handling; EVFA participated in acquisition and analysis of CARS experiments; MC participated in MRI acquisition; JCP participated in acquisition and analysis of neuromuscular junctions; CGB participated in the design of MRI acquisition and analysis; HB participated in acquisition and analysis of CARS experiments; NL participated in the design of the project; YNG participated in the design of the project, performed the experiments in mice and analyzed the data; FEP conceptualized the research, participated in the nonhuman primate experiments, the analysis and data interpretation, wrote the manuscript and final approval. All authors red and approved the final manuscript.

## Acknowledgments

CX3CR1^+/eGFP^ mice were obtained from Dr. Dan Littman (Howard Hughes Medical Institute, Skirball Institute, NYU Medical Centre, New York, USA). Transcriptomic experiments were done at iGE3 Genomics Platform, University of Geneva Switzerland; we thank in particular M. Docquier and C. Delucinge for their assistance in transcriptomic analyses. We thank the Montpellier Rio Imaging facility for confocal acquisitions and the animal facility RAM-CECEMA. We thank H. Noristani for constructive discussions. We also thank P. Villette for his help in video analyses and C. Duperray and H. Hirbec for their help in FACS analysis.

## Availability of supporting data

All data analyzed during this study included in the published article and its supporting information are available from the corresponding author on reasonable request.

