## Supplementary Fig 1, Tab 1 & 2 for "Inhibiting microglia proliferation after spinal cord injury improves recovery in mice and nonhuman primates"

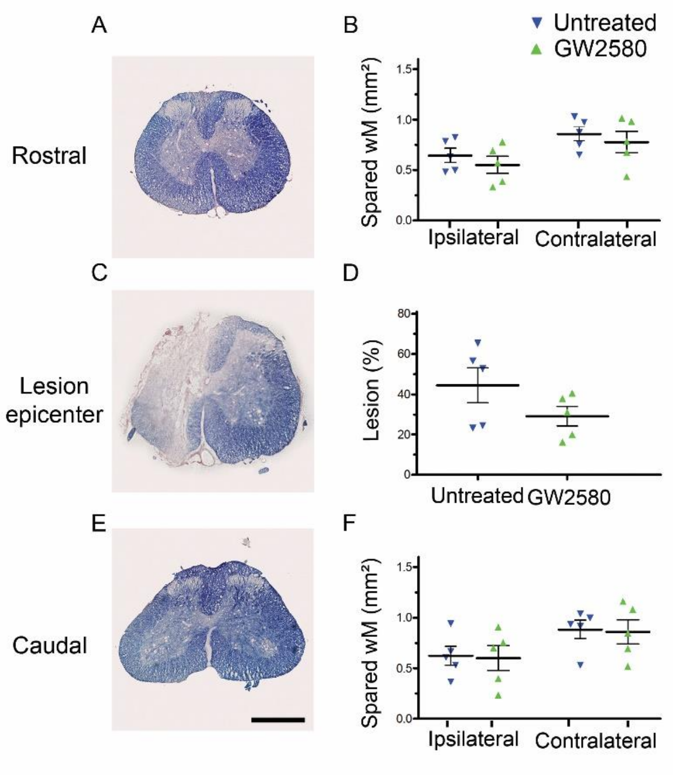


**<u>Supplementary Figure 1:</u> Transient CSF1R blockade after lateral spinal cord hemisection in nonhuman primates does modify lesion extension**

Bright-field micrographs displaying luxol fast blue and neutral red stained axial sections rostral (**A**) within (**C**) and caudal (**E**) to the lesion site at 3 months after SCI in a GW2580-treated nonhuman primate. Luxol-based quantifications of the spared white matter rostral (**B**) and caudal (**F**) to the lesion site as well as the percentage of damaged tissue at the epicenter (**D**).

__Supplementary Table 1.__ **Database of differential expression comparison of microglia from GW2580-treated and untreated mice at 1 week after spinal cord hemisection.** p-value with FDR≤0.05 and FC≥2.

| Gene | Description | GO Process | Fold Change | p-value with FDR |
|---|---|---|---|---|
| ***Cd38*** | CD38 antigen | Negative regulation of neuron projection | -7.71 | 0.0024 |
| ***Cd40*** | CD40 antigen | Immune process | -3.48 | 0.0124 |
| ***Cxcl13*** | Chemokine (C-X-C motif) ligand 13 | Immune process | -3.34 | 0.0214 |
| ***Pf4*** | Platelet factor 4 | Inflammatory response | -3.3 | 0.0155 |
| ***Emilin2*** | Elastin microfibril interfacer 2 | | -3.12 | 0.0057 |
| ***Lyz1*** | Lysozyme 1 | Defense response to bacterium | -2.89 | 0.0402 |
| ***Msr1*** | Macrophage scavenger receptor 1 | Positive regulation of macrophage derived foam cell differentiation | -2.82 | 0.0024 |
| ***Fn1*** | Fibronectin 1 | Positive regulation of cell proliferation regulation | -2.80 | 0.0087 |
| ***Cspg4*** | Chondroitin sulfate proteoglycan 4 | Cell population proliferation, glial cell migration | -2.74 | 0.0107 |
| ***Adm*** | Adrenomedullin | Positive regulation of cell proliferation regulation | -2.59 | 0.0327 |
| ***Cybb*** | Cytochrome b-245. beta polypeptide | Inflammatory response | -2.27 | 0.0024 |
| ***Pltp*** | Phospholipid transfer protein | Lipid metabolic process | -2.27 | 0.0201 |
| ***Gpx3*** | Glutathione peroxidase 3 | Response to oxidative stress | -2.26 | 0.0057 |
| ***Lyz2*** | Lysozyme 2 | Killing of cells of other organism | -2.12 | 0.0134 |
| ***Itsn1*** | Intersectin 1 (SH3 domain protein 1A) | Brain development | -2.07 | 0.0407 |
| ***Gpnmb*** | Glycoprotein (transmembrane) nmb | Positive regulation of cell migration | -2.07 | 0.0051 |
| ***Kcnk12*** | Potassium channel. subfamily K. member 12 | Potassium ion transmembrane transport | 2.06 | 0.0377 |
| ***Srpk3*** | Serine/arginine-rich protein specific kinase 3 | Cell differentiation | 2.08 | 0.0104 |
| ***Sdk1*** | Sidekick cell adhesion molecule 1 | Cell adhesion | 2.32 | 0.0191 |

**<u>Supplementary Table 2.</u> Enrichment analysis in the comparison of microglia from GW2580-treated and untreated mice at 1 week after spinal cord hemisection.** p-value with FDR≤0.05 and FC≥2.

| **Rank** | **Molecular function** | **p-value with FDR** | **Genes in data vs total genes in the pathway** | **Genes** |
|---|---|---|---|---|
| **1** | CXCR3 chemokine receptor binding | 7.314E-4 | 2/5 | *Cxcl13, Pf4* |
| **2** | Heparin binding | 1.150E-3 | 4/221 | *Gpnmb* (*Osteoactivin*), *fibronectin, Cxcl13, Pf4* |
| **3** | Lysozyme activity | 6.219E-19 | 2/13 | *Lyz1, Lyz2* |

| **Rank** | **Processes** | **p-value/ FDR** | **Genes in data vs total genes in the pathway** | **Genes** |
|---|---|---|---|---|
| **1** | Inflammatory response | 5.588E-05 | 8/874 | *Adrenomedullin, Cd40, Cspg4, fibronectin, Cxcl13, Pf4, Cybb, Lyzozym* |
| **2** | Regulation of angiogenesis | 1.487E-04 | 6/484 | *Adrenomedullin, Cd40, Cxcl13, Pf4, Cybb, Gpnmb* (*Osteoactivin*) |
| **3** | Regulation of developmental process | 1.487E-04 | 12/3978 | *Adrenomedullin, Cd38, Cd40, Cspg4, fibronectin, Cxcl13, Cybb, Gpnmb* (*Osteoactivin*), *Gpx, Intersectin, Msr1, Intersectin* |
| **4** | Response to other organism | 1.487E-04 | 10/2427 | *Adrenomedullin, Cd40, fibronectin, Cxcl13, Cybb, Gpx, Gpx3, Lysozyme, Lyz2, Pf4* |
| **5** | Response to external biotic stimulus | 1.487E-04 | 10/2431 | *Adrenomedullin, Cd40, fibronectin, Cxcl13, Cybb, Gpx, Gpx3, Lysozyme, Lyz2, Pf4* |
